# A novel class of conserved sucrose-phosphate phosphatases highlights the diversity of cyanobacterial sucrose metabolism

**DOI:** 10.64898/2026.09.11.751015

**Authors:** María Santos-Merino, Sreeahila Retnadhas, Sigal Lechno-Yossef, Matthew E. Dwyer, Mariana Aubele-Gonzalez, Daniel C. Ducat

**Affiliations:** MSU-DOE Plant Research Laboratory, Michigan State University, East Lansing, MI, United States, 48824; Department of Biochemistry and Molecular Biology, Michigan State University, East Lansing, MI, United States, 48824

**Keywords:** Cyanobacteria, sucrose metabolism, sucrose-phosphate phosphatase (SPP), sucrose-phosphate phosphatase-like (SPP-like), sucrose 6-phosphate, central carbon metabolism

## Abstract

Sucrose metabolism is an important feature of the physiology of the green lineage of photosynthetic organisms and has therefore been the subject of considerable research on plants, algae, and cyanobacteria. Canonical sucrose biosynthesis pathways often involve the condensation of NDP-glucose and fructose-6-phosphate through the action of sucrose-phosphate synthase, then the dephosphorylation of sucrose 6-phosphate into sucrose via sucrose-phosphate phosphatase (SPP). However, many cyanobacterial genomes encode multiple homologs of SPP proteins (SPP-like), including variants that appear to lack key residues reported to be important for sucrose 6-phosphate binding. Herein, we examine these SPP-like proteins, focusing on the biochemical and physiological characterization of the SPP-like protein encoded by the cyanobacterial model, *Synechococcus elongatus* PCC 7942. Bioinformatic analysis suggests that the SPP-like family of proteins is highly conserved across cyanobacterial species and forms distinct phylogenetic clades that are more widely distributed than the canonical SPP proteins themselves. Biochemical analysis of the purified *S. elongatus* PCC 7942 SPP-like protein reveals that it not only retains the capacity to dephosphorylate sucrose 6-phosphate, but it also may have physiologically relevant phosphatase activity on 3-phosphoglycerate (3-PGA). We provide evidence that the SPP-like family of proteins represents a well-conserved group of phosphatases across cyanobacteria and suggest some enzymes in this family may have evolved a broader substrate specificity relative to the well-characterized SPP family. Our results have broader implications for cyanobacterial regulation of sucrose biosynthesis and other key steps of central carbon metabolism.

**Graphical abstract:** 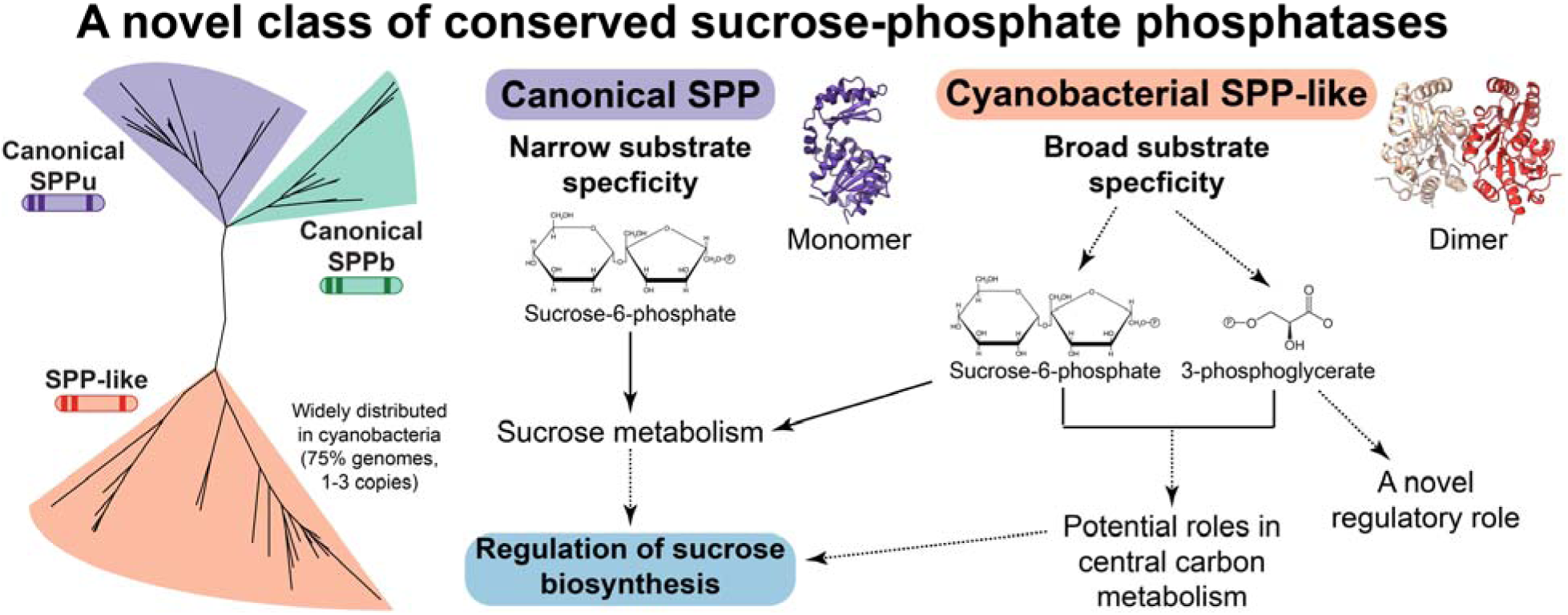

## 1. Introduction

Sucrose synthesis is phylogenetically restricted to oxygenic phototrophs, including cyanobacteria, green algae, and land plants (Salerno & Curatti, 2003), with few reports of sucrose biosynthesis among non-cyanobacterial prokaryotes (MacRae & Lunn, 2012). Sucrose biosynthesis has been proposed to represent an early osmoprotective metabolic pathway in ancient cyanobacteria (Blank, 2013), and was likely inherited by algae and plants through the endosymbiotic acquisition of cyanobacteria that gave rise to chloroplasts (Salerno & Curatti, 2003). Sucrose serves as the primary vehicle for fixed carbon transfer from vegetative cells to heterocysts in filamentous nitrogen-fixing cyanobacterial species (Curatti et al., 2002), similar to its role as a carbon and energy carrier in plants and algae. More specifically, sucrose is the most important metabolite involved in resource allocation and energy balance, since it is the major end product of photosynthetic carbon metabolism and the predominant form of carbon transported to heterotrophic tissues in most plants (Lepper et al., 2025). Finally, sucrose is being widely explored as a cyanobacteria-derived carbohydrate feedstock for both direct and co-culture bioproduction systems (Santos-Merino et al., 2023).

The majority of cyanobacteria synthesize sucrose using the same synthesis pathway as plants, involving the sequential action of two enzymes: sucrose-phosphate synthase (SPS) and sucrose-phosphate phosphatase (SPP) (Santos-Merino et al., 2023). Additionally, some cyanobacterial species synthesize sucrose via an alternative, bidirectional enzyme, sucrose synthase (SuS) (Curatti et al., 2008), an enzyme well-characterized in plants and cyanobacteria to regulate directionality by both substrate availability and pH (Porchia et al., 1999; Schmolzer et al., 2016).

Notably, the enzymes involved in sucrose biosynthesis adopt a strikingly modular architecture (Salerno & Curatti, 2003). Specifically, SPS and SPP are frequently encoded either as unidomainal enzymes, or as a bidomainal fusion of SPS and SPP domains connected by a peptide linker. Moreover, genes encoding SPP function appear to have undergone frequent endoduplication over evolutionary timescales, giving rise to many species of cyanobacteria and plants with multiple copies of SPP genes. As one illustrative example, the genome of the cyanobacterial model *Synechocystis* sp. PCC 6803 encodes a bidomainal SPS-SPP (confusingly formally named SPS) where the SPP domain contains mutations in key catalytic residues that render it enzymatically dead (Fieulaine et al., 2005). Apparently to compensate for the lack of SPP activity of this bidomainal SPS, *Synechocystis* sp. PCC 6803 encodes a separate, active unidomainal SPP in its genome (Lunn, 2002).

Further illustrating the modularity of sucrose biosynthesis genes, the existence of another family of so-called “SPP-like” proteins has recently been reported for cyanobacteria (Bai et al., 2025; Santos-Merino et al., 2023). These SPP-like proteins are phylogenetically distinct from the canonical SPP uni-or bi-domainal proteins and contain mutations in key residues involved in binding to sucrose 6-phosphate (S6P) (Santos-Merino et al., 2023). The presence of these mutations potentially suggests that SPP-like proteins lack the ability to dephosphorylate S6P. Additionally, the SPP-like family appears to be widely conserved across the cyanobacterial phylum with some species containing more than two genes encoding for different SPP-like proteins in addition to other canonical SPP proteins (Santos-Merino et al., 2023). SPP-like proteins are frequently annotated in genomic databases as (putative) members of the haloacid dehydrogenase superfamily (HAD-superfamily) hydrolases subfamily IIB, and they contain three conserved HAD motifs associated with the active site in canonical SPPs (Fieulaine et al., 2005).

Notably, plants also exhibit SPP proteins that appear to lack enzymatic activity, and such inactive SPPs have been hypothesized to have noncatalytic moonlighting functions as transcriptional regulators (Albi et al., 2016). By analogy, catalytically-dead plant homologs of trehalose-phosphate phosphatases (TPPs) and TPP-like proteins have been shown to perform non-catalytic moonlighting functions as metabolite sensors or as transcriptional regulators, regulating plant signaling mediated by trehalose 6-phosphate (T6P) (Fichtner, 2025). However, any possible signaling roles for SPP proteins - or S6P itself - are not well-supported by experimental evidence and therefore remain speculative (Lunn & MacRae, 2003; Chen et al., 2005; Chen et al., 2008).

There is a major research gap in understanding the role of SPP-like proteins in cyanobacteria. Preliminary analysis has suggested that this family of enzymes is likely unable to utilize S6P based on the lack of conservation of residues involved in S6P binding (Santos-Merino et al., 2023). This could suggest that SPP-like enzymes are involved in dephosphorylation of alternative substrates; however, this has not been experimentally explored. The present work reports a detailed biochemical characterization of the SPP-like protein from *Synechococcus elongatus* PCC 7942 (Synpcc7942_0566; SPP-like_7942_). In comparison to the well-studied canonical SPP from *Synechocystis* sp. PCC 6803 (SPP_6803_; Slr0953), SPP-like_7942_ exhibited a broader substrate specificity and greater tolerance to acidic pHs in comparison to SPP_6803_. Surprisingly, despite the loss of key residues reported to bind S6P within the active site, SPP-like_7942_ exhibited an apparent comparable to that of SPP_6803_, although SPP_6803_ displayed a markedly higher turnover rate (*K_cat_*) relative to SPP-like_7942_. Taken together, our results suggest that SPP-like_7942_ and the canonical SPP are simultaneously conserved in *S. elongatus* PCC 7942, perhaps due to differential regulation or non-overlapping functions of these two proteins.

## Materials and methods

### 1.1. Sequence homology

The protein sequences of homologs of SPP-like_7942_, SPSu (unidomainal SPS), SPPb (SPP domain of the bidomainal SPS-SPP), and SPPu (unidomainal SPP) were obtained from the NCBI and UniProt databases. Sequences were retrieved using BLAST tools in both databases; established cyanobacterial enzymes in these pathways were used as queries: SPSu from *Nostoc* sp. PCC 7120 (GenBank accession No. BAB76075.1) (Cumino et al., 2002), SPSb from *S. elongatus* PCC 7942 (GenBank accession No. ABB56840.1) (Liang et al., 2020), SPPu from *Synechocystis* sp. PCC 6803 (SPP_6803_; GenBank accession No. BAA18419.1) (Fieulaine et al., 2005) and SPP-like from *S. elongatus* PCC 7942 (SPP-like_7942_; GenBank accession No. ABB56598.1) (Santos-Merino et al., 2023). Homologs were retrieved for each protein across 119 cyanobacterial genomes, retaining hits with E-values ≤10^−15^, identity ≥35% and coverage ≥80%. The retrieved sequences are listed in Table S1.

### 1.2. Multiple sequence alignments and phylogenetic trees

Multiple sequence alignments (MSAs) for SPP-like, SPPu, SPPb and HAD-hydrolase type IIB proteins were generated using MAFFT with the G-INS-I algorithm. Each alignment was trimmed using AlignmentViewer to remove positions containing ≥85% gaps. Abnormally short and phylogenetically distant sequences were manually removed. Sequence logos of conserved motifs for each analyzed enzyme were obtained using WebLogo server. Maximum-likelihood phylogenetic trees were inferred using IQ-TREE with automatic model selection performed by ModelFinder. The LG substitution model was selected, and branch support was assessed using 1,000 ultrafast bootstrap replicates and 1,000 SH-aLRT replicates (-alrt 1000 -bb 1000). Tree visualizations were generated using FigTree. ConSurf was used to find structural and functional residues of enzymes and conserved amino acid positions.

### 1.3. Molecular cloning, protein expression, and purification

Full-length SPP_6803_ and SPP-like_7942_ were cloned into a pET28a vector (Novagen, Darmstadt, Germany), with an N-terminal 6xHisTag (pSL501 and pSL502, respectively). SPP_6803_ and SPP-like_7942_ were expressed in *Escherichia coli* strain BL21 (DE3) (Invitrogen) by induction of cultures with 50 μM IPTG at OD_600_ = 0.4–0.7 at 37 °C, and induced cultures were incubated for 18–24 h at 18 °C. Cultures were harvested by centrifugation and pellets were stored at −20 °C until purification.

For His-tagged protein purification, cell pellets were resuspended in lysis buffer (25 mM HEPES, 150 mM KCl, 5 mM MgCl_2_ and 10% glycerol) containing EDTA-free SigmaFast protease inhibitor cocktail (Sigma, St. Louis) and DNaseI was added to a final concentration of approximately 50 µg mL^−1^. Cells were lysed by two passes through a French Press at 16,000 psi cell pressure. Cell lysates were clarified by centrifugation at 45,000 ×g for 30 min, and the clear lysate was passed through a 0.45 µm filter. 50 mM imidazole was added to filtered lysates which were loaded on a HisTrap column (GE Healthcare, Little Chalfont, UK), equilibrated with lysis buffer containing 50 mM imidazole, attached to an AKTA Pure FPLC (GE Healthcare, Little Chalfont, UK). The column was washed with 10 CV of loading buffer, then bound proteins were eluted with a gradient of 50 mM to 500 mM imidazole in the same buffer over 10 CV. Proteins were observed on SDS-PAGE gels stained with Coomassie blue. Protein concentration was quantified using the BCA assay kit (Sigma) against a bovine serum albumin standard.

### 1.4. Molecular mass determination

The apparent molecular masses of SPP_6803_ and SPP-like_7942_ were estimated by analytical size-exclusion chromatography using Superdex® 200 Increase 10/300 GL column at a 0.5 mL min^−1^ flow rate. The column was calibrated with the standard protein molecular weight marker (151-1901, Bio-Rad, USA) containing thyroglobulin (670 kDa), γ-globulin (158 kDa), ovalbumin (44 kDa), myoglobin (17 kDa), and vitamin B12 (1.35 kDa). Bovine serum albumin (BSA; 66 kDa) (B14, Thermo Fisher Scientific, USA) was added as an additional standard. Apparent molecular masses were estimated from a standard curve of the logarithm of the molecular mass versus the ratio of the elution volume (Ve) to the column void volume (Vo). The oligomeric states of both proteins were further assessed by Blue-native gel electrophoresis (Text S1) and structural predictions generated with AlphaFold2 Multimer (Text S2).

### 1.5. Phosphatase activity assays

Initial activity assays for SPP_6803_ and SPP-like_7942_ utilized the universal substrate pNPP. The substrate hydrolysis was monitored at RT (room temperature) using the Cary 60 UV-Vis spectrophotometer (Agilent Technologies, CA, USA). A 500 µL assay reaction mixture containing 4 mM pNPP, 5 mM MgCl_2_, 32 µg enzyme, at the indicated pH: 10 mM acetate buffer for pH 3.5 – 5.0, 50 mM MES buffer for pH 6.0, and 50 mM HEPES buffer for pH 7.0 – 8.0. Reaction mixtures were monitored for increased absorbance at 405 nm over 10 min as a measure of *p*-nitrophenol product formation, using at least three independent reactions were measured, and the blank signal was subtracted. Protein stability was measured at each indicated pH after 30 min incubation.

Similarly, steady-state kinetic assays were performed in a 500 µL assay reaction mixture containing 5 mM MgCl_2_, 32 µg enzyme, 50 mM HEPES buffer pH 7.0 for SPP_6803_ or 10 mM acetate buffer pH 4.0 for SPP-like_7942_ and different concentrations of pNPP (0.2 – 5 mM). At least three readings were recorded for each pH, and the blank-subtracted average was utilized for statistical analysis.

Phosphatase activities of SPP_6803_ and SPP-like_7942_ against putative phosphorylated metabolites were determined by measuring phosphate released using the PiPer™ Phosphate Assay Kit (Life Technologies, USA); a colorimetric assay based on a reaction output converting Amplex Red Reagent (10-acetyl-3,7-dihydrophenoxazine) to resorufin, which has absorption/emission maxima of 563 nm and 587 nm, respectively. Assays were conducted in a 100 μL mixture (5 mM MgCl_2_, 50 mM HEPES buffer pH 7.0 for SPP_6803_ and for SPP-like_7942_ or 10 mM acetate buffer pH 4.0 for SPP-like_7942_ and different concentrations of each substrate). First, enzyme was titrated to keep the conversion of substrate during the 10 min reaction to <10% of the total substrate used. For SPP_6803_ assays, these concentrations were determined to be 3 µg and 0.02 µg for G6P and S6P, respectively. For SPP-like_7942_, 0.1 µg, 6 µg and 5 µg for G3P, 3PGA and S6P were used, respectively. 50 μL phosphatase reaction mixture was combined with 50 μL PiPer™ reaction reagent, incubated at RT for 10 min before measuring the fluorescence using the SpectraMax® iD3 microplate reader (Molecular Devices, San Jose, CA, USA; excitation at 530 nm and emission at 590 nm). Measurements from paired blanks lacking substrate were subtracted from sample values before calculating steady-state kinetic parameters.

Protein phosphatase activity was also determined using the procedures described in Text S3.

### 1.6. CD spectroscopy

CD spectra were collected on a Chirascan plus Spectrometer (Applied Photophysics, Leatherhead, UK) at RT using 1 mm path length quartz cuvettes. Protein samples of SPP_6803_ and SPP-like_7942_ were diluted to 10 µM in 10 mM potassium phosphate buffer adjusted to different pH, incubated on ice for 10 min before measuring CD spectra. The denaturation process was tracked by collecting kinetic (50 ms) scans between 190 – 280 nm with 1 nm bandwidth. Traces represent averages of four consecutive scans after buffer background subtraction. The CD spectral scans were smoothened using OriginPro 2024 (OriginLab Corporation, USA) and were converted to mean residues ellipticity using the formula: [θ] = θ_obs_·MRW/10·l·c, where [θ] is the molar ellipticity, θ_obs_ is the measured ellipticity in millidegrees, MRW is the mean residual weight of the protein (123 for SPP_6803_ and 118 for SPP-like_7942_), l is the cell path length in centimeters, c is the protein concentration in gr L^−1^.

### 1.7. Pull-down assays

Purified His□-tagged SPP-like_7942_ was used as bait to pull down potential interacting partners from total protein extracts of *S. elongatus* PCC 7942 obtained by bead-beating in 150 µL lysis buffer (0.3 mM EDTA, 1 mM DTT, 25 mM Tris pH 7.4, 50 mM NaCl, and protease inhibitor). Cells were disrupted using a TissueLyser (Qiagen, Germantown, MD) with glass beads (0.1 mm; Biospec Products, Bartlesville, OK) at a frequency of 30 Hz, with 30 cycles of 1 min on and 1 min off on ice. The lysate and beads were pelleted at 2,000 ×g for 10 min at 4□°C and supernatant was collected and cleared via centrifugation at 12,000□×g for 10□min at 4□°C. Supernatant was transferred to a new tube and flash frozen.

Purified His□-tagged SPP-like (90 µg in 100 µL HEPES buffer) was loaded onto a Ni²□-NTA spin column (15 µL of 50% Ni-NTA agarose suspension, 30210, Qiagen, Hilden, Germany) pre-equilibrated with HEPES buffer (25 mM HEPES, 150 mM KCl; 5 mM MgCl_2_ and 10% glycerol). The column was incubated at 4 °C for 30 min with gentle rocking to allow immobilization of the bait protein, then bottom cap was removed, and the column was centrifuged at 1,000 rpm for 1 min to remove unbound protein. Cyanobacterial cell lysate (500 µg of total protein diluted in 250 µL HEPES buffer) was added to the Ni²□-NTA bound SPP-like_7942_ and incubated overnight at 4 °C to allow interaction with potential binding partners. Excess lysate was removed by centrifuging the column at 1,000 rpm for 1 min and the column was washed three times with 30 µL HEPES buffer containing 50 mM imidazole. Bound prey proteins were eluted using 20 µL HEPES buffer supplemented with 300 mM imidazole and collected by centrifugation. As a negative control, parallel pull-down experiments were performed using Ni²□-NTA resin without immobilized bait protein.

### 1.8. In-gel protein digestion and extraction

Proteins recovered via pull-down were separated on Any kD Mini-PROTEAN TGX precast gels (Bio-Rad, Hercules, CA) and stained with Coomassie Brilliant Blue R-250 (Thermo Fisher Scientific, Somerset, NJ). Lanes for each replicate (n = 2) were manually cut. Each band was destained with 10% methanol/5% glacial acetic acid (v/v). Prior to protein identification by mass spectrometry, in-gel digestion was performed following the protocol described in Text S4.

### 1.9. LC–MS/MS analysis and LC–MS/MS data processing

LC-MS/MS peptide analysis was performed using an EASY-nLC1200 UHPLC (Thermo Fisher Scientific, Waltham, MA, USA) interfaced via nanoSpray Flex ion source to an Orbitrap Fusion Lumos Tribrid MS (Thermo Fisher Scientific, Waltham, MA, USA). Protein quantification was achieved using the label-free quantification (LFQ) algorithm in MaxQuant and protein ratios were determined by the LFQ intensity. A detailed description of the LC-MS/MS analysis and data processing workflow is provided in Text S5.

### 1.10. Bacterial two hybrid and β-galactosidase activity assay

N-terminal and C-terminal fusions of the T25 domain to SPP-like_7942_ were constructed using the plasmids pKTN25 and pKT25, respectively, whereas T18 fusions to selected pull-down candidates were built in plasmids pUT18 and pUT18C, respectively. All plasmids were sequence-verified and co-transformed into *E. coli* BTH101 in pairwise combinations. Colonies of T18/T25 cotransformants were cultured overnight in LB medium supplemented with 50 μg mL^−1^ carbenicillin, 50 μg mL^−1^ kanamycin, and 0.5 mM IPTG at 30 °C with shaking at 250 rpm. Overnight cultures were spotted onto LB agar plates containing X-gal supplemented with 50 μg mL^−1^ carbenicillin, 50 μg mL^−1^ kanamycin, and 0.5 mM IPTG. Plates were incubated in the dark at 30°C for at least 24 h before imaging. β-galactosidase activity was measured to quantify protein-protein interactions. 100 µL of each overnight culture were transferred to an individual well of a 96-well deep-well plate containing 1 mL of Z buffer (60 mM Na_2_HPO_4_, 40 mM NaH_2_PO_4_, 10 mM KCl, 1 mM MgSO_4_, 38.5 mM β-mercaptoethanol). Subsequently, 20 µL of 0.01% SDS and 50 µL of chloroform were added to each well, and the samples were mixed by pipetting up and down five times to permeabilize the cells. After allowing the chloroform to settle for 2 min, 100 μL of the aqueous phase were transferred to a flat-bottom 96-well microplate. The reaction was initiated by adding 40 μL of o-nitrophenyl-β-D-galactopyranoside (ONPG, 2 mg mL^−1^). As yellow color developed, the reaction was stopped by adding 50 µL of 1 M Na_2_CO_3_. β-galactosidase activity was expressed as Miller Units and calculated according to the following equation: Miller Units = 1000× [(Abs_420_-(1.75×Abs_550_))]/(T×V×Abs_600_), where Abs_420_ and Abs_550_ were measured from the reaction mixture, Abs_600_ was measured from the overnight culture, T is the reaction time (min) of before adding 1 M Na_2_CO_3_, and V is the volume of culture used in the assay (mL). All absorbance measurements were performed using a SpectraMax® iD3 microplate reader (Molecular Devices, San Jose, CA, USA).

### 1.11. Gene neighborhood analysis

To analyze the genomic context of SPP-like sequences, representative protein sequences were submitted to EFI-GNT, and genes located within ±10 open reading frames (ORFs) were examined. Gene neighborhood diagrams were annotated based on the EFI-generated output diagrams and UniProt functional annotations.

### 1.12. Strains, culture conditions and strain construction

Detailed descriptions of the culture conditions used for the experiments involving cyanobacterial strains are provided in Text S6. All strains used in this study are listed in Table S2. Strain construction was performed as described in Text S7 using the plasmids listed in Table S3.

### 1.13. Photosynthetic fluorescence measurements

Measurements of apparent quantum yield of PSII (Φ_II_) were performed on a custom-built fluorimeter/spectrophotometer as described previously. See Text S8 for a detailed protocol.

### 1.14. Microscopy and image analysis

Live-cell microscopy was performed using exponentially growing cells as previously described (Santos-Merino et al., 2025). A detailed microscopy protocol and image analysis workflow are provided in Text S9.

## 2. Results and discussion

### 2.1. SPP-like is widely conserved in cyanobacteria and phylogenetically distant from canonical SPP

SPP-like proteins are highly conserved across the phylum cyanobacteria, even in species that encode one or more canonical SPP genes (Figs. 1A and S1 and Table S1). Cyanobacterial SPP-like proteins cluster into at least three groups that are distantly related to the canonical SPPu and SPPb domains, whereas plant SPP homologs are phylogenetically closely related to cyanobacterial SPPu proteins (Figs. 1A and S1). Remarkably, our phylogenetic survey suggested that members of the SPP-like gene family are more widespread than canonical SPP genes, with many genomes encoding two or more SPP-like proteins (Figs. 1B and S2). We found many cyanobacterial genomes encode SPP-like homologs while lacking canonical SPPu or SPPb genes (Figs. 1B and S2A), providing indirect evidence that the SPP-like protein family may play distinct roles relative to the better-characterized functions of SPP in sucrose metabolism. We expanded our analysis to correlate the presence of SPP-like genes and cyanobacterial habitat (Figs. 1B and S2A). Broadly, cyanobacteria associated with freshwater and terrestrial environments encoded more copies of SPP-like genes, while ∼23.5% of genomes lacking a gene encoding SPP-like were mostly originating from marine habitats. The highest prevalence of SPP-like genes was found in filamentous cyanobacteria, particularly heterocyst-forming species, which often encode multiple SPP-like homologs (Fig. S2B). Only five species lacked any version of SPP (SPPu, SPPb and SPP-like), and at least two of these species (*Arthrospira* spp.) encode SuS as the likely pathway for sucrose synthesis.

**Fig. 1.**
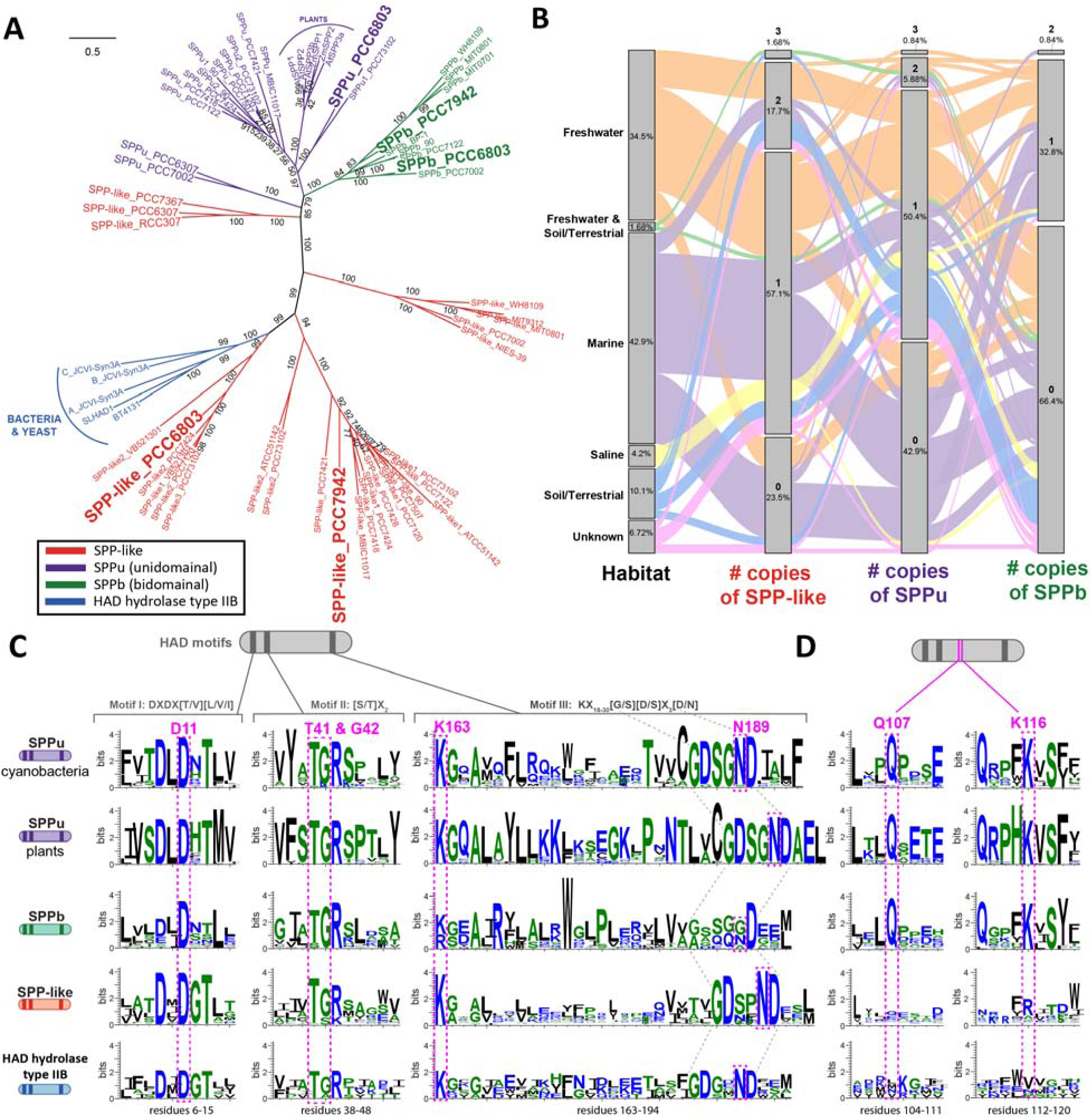
Diversity of SPP and SPP-like proteins across different groups. **(A)** Phylogenetic tree showing the evolutionary relationships between SPP unidomainal (SPPu; purple), SPP bidomainal (SPPb; green) and SPP-like (red) of a subset of cyanobacteria and plants, as well as HAD hydrolase type IIB from bacteria and yeast (blue). A more extended version of this tree can be found in Fig. S1. Numbers at each branch point indicate the bootstrap support values (percentages) calculated from 1,000 replicate trees using IQ-TREE. **(B)** Alluvial map depicting the relationship between the habitat of 119 cyanobacterial species and the number of copies of SPP-like and SPP identified in their genomes. Strains and accession numbers of each gene can be found in Table S1. **(C)** Sequence logos for conserved motifs in phosphatases, with key residues involved in S6P binding highlighted in pink. **(D)** Sequence logos for conserved residues binding to S6P located outside the phosphatase motifs. **(C, D)** These residues were identified from the crystal structure of SPP_6803_ (Fieulaine et al., 2005).

SPP and SPP-like homologs belong to the larger superfamily of HAD-hydrolases, in the subfamily IIB (Albi et al., 2016; Santos-Merino et al., 2023). The HAD superfamily comprises mainly uncharacterized enzymes and is named after the first structurally characterized family member, 2-haloacid dehalogenase (Hisano et al., 1996). Most known catalytic activities carried out by HAD family members involve phosphoryl transfer reactions, with only a few members shown to exhibit phosphatase activity: beta-phosphoglucomutase, phosphonatase, or dehalogenase activities (Burroughs et al., 2006). Only a limited number of HAD hydrolases belonging to the subfamily IIB have been studied, all from non-photosynthetic bacteria (Haas et al., 2022; Kaur et al., 2024; Lu et al., 2005). Interestingly, the previously characterized bacterial HAD hydrolases from subfamily IIB are phylogenetically adjacent to the largest clusters of SPP-like cyanobacterial proteins, including the SPP-like homolog of *S. elongatus* PCC 7942 (Figs. 1A and S1). However, only a limited number of these enzymes display measurable activity toward sugar phosphates, with unknown physiological significance. Indeed, the catalytic activity of the protein BT4131 from *Bacteroides thetaiotaomicron* VPI-5482 showed very limited activity against S6P, suggesting that other metabolites are likely to represent its physiologically relevant substrates (Lu et al., 2005).

The HAD superfamily is characterized by three conserved motifs (I, II and III) that are thought to be critical for phosphatase and phosphotransferase reactions (Fieulaine et al., 2005) (Figs. 1C, S3A and S3B). The key catalytic residues within motifs I, II, and III are highly conserved across SPPu, SPP-like, and HAD hydrolase type IIB members analyzed (Fig. 1C), with the exception of certain members of the SPPb family, which have been shown to lack catalytic activity (Liang et al., 2020; Lunn et al., 2003). By contrast, other conserved residues that have been proposed to bind to S6P (and therefore confer substrate specificity) are poorly conserved in SPP-like and HAD hydrolase type IIB members (Fig. 1D). Specifically, the crystal structure of SPPu from the cyanobacterium *Synechocystis* sp. PCC 6803 was used to determine residues involved in binding to S6P: Asp11, Thr41, Gly42, Gln107, Lys116, Lys163 and Asn189 (Fieulaine et al., 2005) (Figs. 1C and 1D). As mentioned previously, some SPPb homologs lack catalytic activity, and mutations in Asp11, Lys163, and/or Asn189 are associated with loss of enzymatic function (Liang et al., 2020; Lunn et al., 2003), while other HAD family members exhibit high conservation of these residues (Liang et al., 2020; Lunn et al., 2003). Among the remaining S6P-binding residues, only Gln107 and Lys116 (located between motifs II and III; Fig. 1D) are not strictly conserved, exhibiting substantial variability in both SPP-like proteins and representative HAD hydrolase subfamily IIB members.

Given the variability of key residues implicated in S6P binding and the low sequence similarity among these proteins (Fig. S3), we sought to further characterize the SPP-like protein family to identify potential alternative functions relative to canonical SPP enzymes. The SPP-like protein from *S. elongatus* PCC 7942 (Synpcc7942_0566; SPP-like_7942_) was selected for further study and compared to the relatively well-characterized unidomainal SPP from *Synechocystis* sp. PCC 6803 (SPP_6803_), based on their low sequence similarity but high structural similarity (Fig. S3).

### 2.2. SPP-like_7942_ is a dimer in solution

SPP_6803_ and SPP-like_7942_ were heterologously expressed in *E. coli.* Both proteins predominantly partitioned to the soluble fraction (Fig. 2A), where they could be purified via a single step of Ni²□-affinity chromatography. In SDS-PAGE analysis, both SPP_6803_ and SPP-like_7942_ migrated to positions consistent with their expected molecular weights of 29.9 and 29.46 kDa, respectively (Fig. 2A). Size-exclusion chromatography (SEC) revealed that SPP-like_7942_ eluted at an apparent dimeric size of ∼52 kDa, consistent with structural predictions from AlphaFold, which indicate that the protein forms dimers through interactions between its core domains with high confidence (Fig. S4A and S4B). Native PAGE analysis indicated that SPP-like_7942_ migrated as a single band, while SPP_6803_ migrated as two fractions (Fig. S4C), which were estimated to correspond to ∼17 kDa and ∼40 kDa via SEC (Fig. 2B). Although many bacterial HAD phosphatases are monomeric (Bang et al., 2024; Kawamura et al., 2008; Lu et al., 2005), several bacterial members of this superfamily have been reported to form dimers (Goncalves et al., 2011; Kaur et al., 2024; Suthisawat et al., 2020), indicating that oligomerization among bacterial HAD phosphatases is not uncommon. However, we note that, under alternative experimental conditions, other studies have found SPP_6803_ predominantly in a monomeric form in solution by SEC (Lunn, 2002) and to crystallize as a monomer (Fieulaine et al., 2005). Taken together, our data are consistent with SPP-like_7942_ forming a dimer in solution, which contrasts with the likely monomeric oligomerization state of SPP_6803_, but which may have superficial similarities to plant SPPs, which dimerize through a non-conserved C-terminal domain (Albi et al., 2016; Lunn, 2002).

**Fig. 2.**
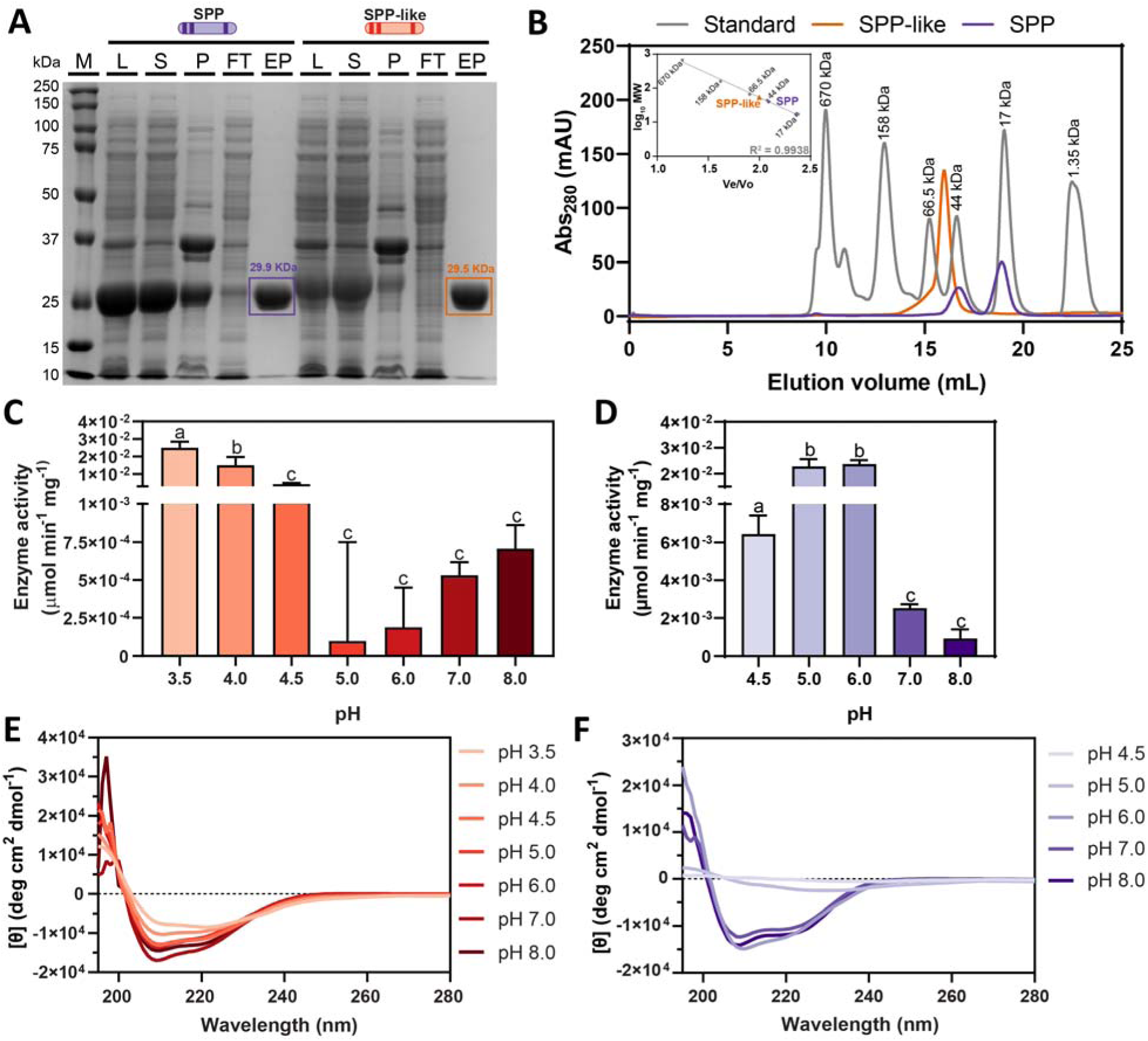
Purification, molecular size determination, and effects of pH on the enzyme activity and secondary structure of SPP_6803_ and SPP-like_7942_. **(A)** SDS-PAGE showing SPP_6803_ and SPP-like_7942_ purified by Ni^2+^-NTA chromatography. M, protein marker; L, lysate; S, supernatant; P, pellet; FT, flow-through; EP, elution peak. **(B)** Size-exclusion chromatography using a Superdex® 200 Increase 10/300 GL column compared to standards monitored at Abs_280_. The calibration curve is displayed on the inset. **(C)** Enzymatic activity of SPP-like_7942_ at different pH using pNPP as the substrate. **(D)** Enzymatic activity of SPP_6803_ at different pH using pNPP as the substrate. **(E)** Far-UV circular-dichroism (CD) spectra of SPP-like_7942_ when subjected to different pH. **(F)** Far-UV circular-dichroism (CD) spectra of SPP_6803_ when subjected to different pH. Negative ellipticity bands at 208 nm and 222 nm are characteristic of α-helical structure. **(C, D)** Averages of ≥3 independent measurements are shown + SD. Significance was calculated by one-way ANOVA followed by Tukey’s multiple comparison test. Bars labeled with different letters are significantly different (P < 0.05). **(E, F)** [θ] is the molar ellipticity. Averages of ≥3 independent measurements are shown.

### 2.3. SPP-like_7942_ and SPP_6803_ exhibit phosphatase activities towards pNPP

To evaluate phosphatase activities of SPP_6803_ and SPP-like_7942_, we first used the universal phosphatase substrate, pNPP (*p*-nitrophenyl phosphate), a substrate that yields *p*-nitrophenol when dephosphorylated, which possesses an absorbance peak at 405 nm (Tietz et al., 1983). At the physiologically relevant pH of 7.0, SPP-like_7942_ showed minimal activity (Fig. 2C), whereas SPP_6803_ exhibited detectable pNPP phosphatase activity (Fig. 2D). We therefore screened both enzymes across a range of pH values, observing clear promotion of SPP-like_7942_ activity under acidic conditions (Fig. 2C). SPP-like_7942_ exhibited a bimodal pH-response profile, potentially implying complex pH-dependent regulation of catalytic activity. SPP_6803_ also exhibited enhanced activity at lower pH (Fig. 2D), but SPP_6803_ activity decreased over time when assayed under acidic conditions (Fig. S5A). The optimal activity of SPP_6803_ at near-neutral pH is consistent with multiple reports for plant SPPs, which show pH optima between 6.0 and 7.0 (Echeverría & Salerno, 1993; Echeverría & Salerno, 1994; Hawker & Hatch, 1966). By contrast, SPP-like_7942_ activity was maintained over time under acidic conditions as low as pH 3.5-4.0 (Figs. 2C and S5B).

In HAD phosphatases, a conserved active-site aspartate (generally denoted as Asp+2) acts as a general acid/base, which is protonated under acidic conditions, enabling it to donate a proton to the leaving group. This protonation stabilizes the developing negative charge during phosphoryl transfer, facilitating cleavage of the phosphate-ester bond (Gohla, 2019). Therefore, as reported with other HAD enzymes (Kaur et al., 2024), reduced pH can sometimes improve catalysis rates *in vitro*. By contrast, the gradual reduction in enzyme activity at lower pH is frequently characteristic of protein denaturation, a possibility we examined through the use of circular dichroism (CD) spectroscopy. Consistent with this interpretation, incubation of purified SPP_6803_ at pH < 6.0 led to the loss of spectral features of secondary structure, while SPP-like_7942_ maintained signatures of secondary structure down to pH 3.5 (Figs. 2E and 2F).

We next determined the kinetic behavior of both SPP-like_7942_ and SPP_6803_ using pNPP as substrate at pH 4.0 and pH 7.0, respectively (Fig. S6). SPP-like_7942_ displayed a sigmoidal plot of the initial reaction rate versus pNPP concentration, rather than the expected hyperbolic plot observed for SPP_6803_. The sigmoidal nature of the SPP-like_7942_ kinetics and the extremely high Hill coefficient (∼12) are inconsistent with a mechanistic interpretation based on classical cooperativity (Holt & Ackers, 2009). We interpret the unusually high Hill coefficient of SPP-like_7942_ as likely indicative of the use of a non-physiological substrate relative to the kinetics typically observed with genuine metabolic substrates (Mercan & Bennett, 2010). Indeed, similar behavior has been previously reported for other HAD enzymes, which display different kinetic plots when different substrates were tested (Kuznetsova et al., 2015). Taken together, the increased activity of SPP-like_7942_ toward pNPP at lower pH is likely a consequence of the greater structural stability of SPP-like_7942_ under acidic conditions, together with the protonation state of key active-site residues in vitro, rather than a physiologically-relevant activation mechanism. Further structural and mechanistic studies will be required to determine the basis of this pH-dependent behavior.

### 2.4. Identification of potential substrates of SPP-like_7942_ by establishing its related metabolic pathway

True metabolic substrates of HAD enzymes are difficult to predict based solely on phylogeny or sequence conservation (Kuznetsova et al., 2006). Therefore, we conducted additional analyses to narrow down potential endogenous targets of SPP-like_7942_. First, we used comparative genome analysis to infer potential functional associations through gene linkage (Fischbach & Voigt, 2010), however, no strong conservation patterns were evident (Fig. S7), consistent with previous observations for HAD family members (Haas et al., 2022). We therefore performed pull-down experiments using SPP-like_7942_ as bait, identifying a total of 100 proteins that were enriched in one or two replicate experiments relative to controls and that could illuminate potential pathways related to the function of SPP-like_7942_ (Fig. 3A and Table S4). Most identified proteins were associated with energy production and conversion, translation, ribosomal structures and biogenesis, and with general/unknown function (Fig. 3B). However, the co-precipitated proteins also included components of metabolic pathways associated with the metabolism of lipids, nucleotides, and amino acids as well as several proteins directly related to photosynthesis. Additionally, a high percentage of co-precipitated proteins were predicted to a cytosolic localization (Table S5). This observation was consistent with an independent analysis of reporter strains expressing fluorophore-tagged SPP-like_7942_ which demonstrated clear cytosolic localization in *S. elongatus* PCC 7942 (Fig. S8), similar to the cytosolic localization previously reported for plant SPP proteins (Echeverría & Salerno, 1993).

**Fig. 3.**
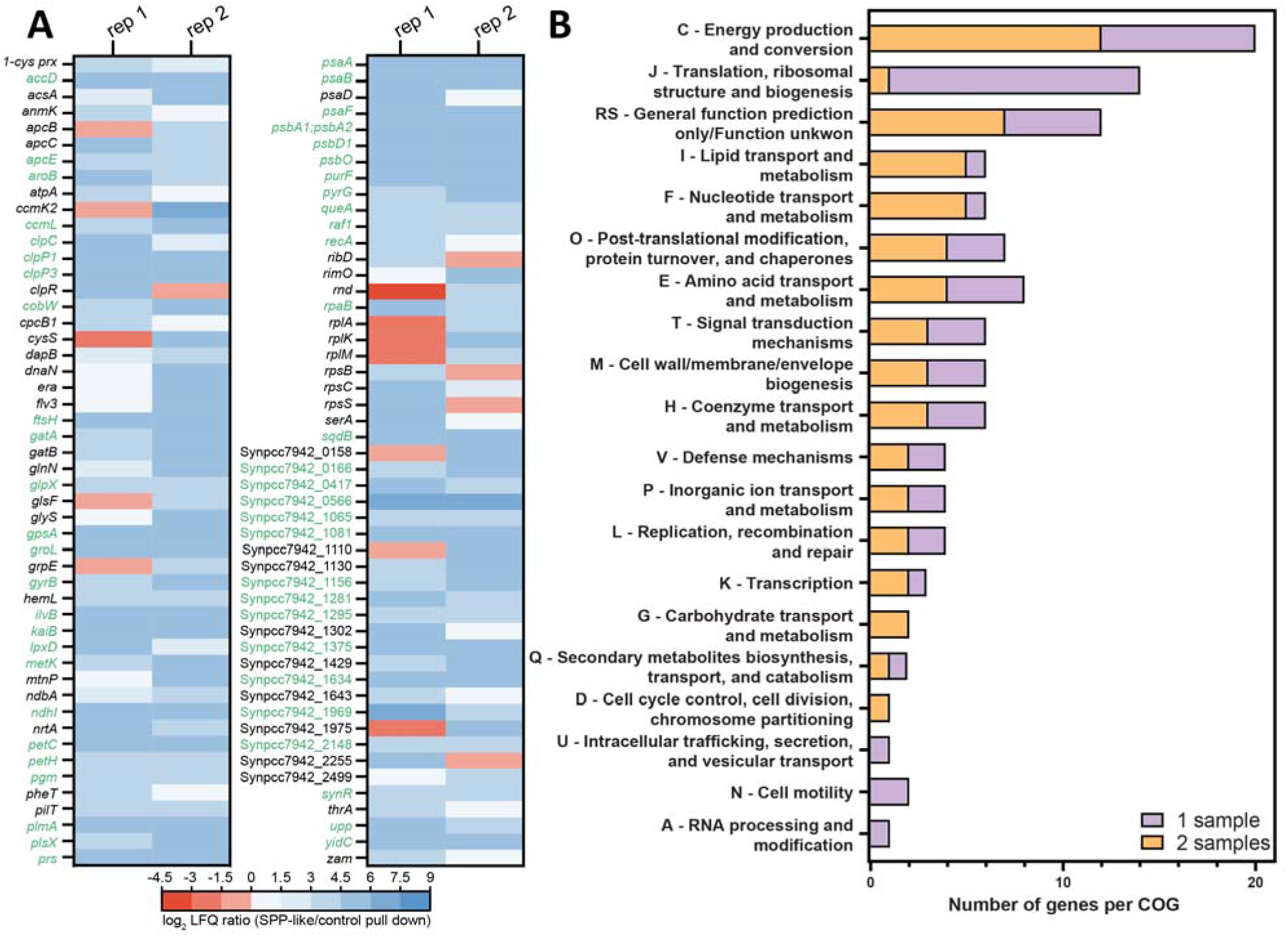
Proteomic analysis of the SPP-like_7942_ interactome. **(A)** Heatmap of the SPP-like_7942_ interactome identified using Ni² -NTA-immobilized His-tagged SPP-like_7942_ as bait. Log - transformed LFQ ratios of proteins co-precipitated with SPP-like_7942_ relative to the empty-bead control are displayed. Proteins identified as hits in at least two replicates are denoted in green, whereas proteins identified in one replicate are indicated in black. **(B)** Functional category composition of putative interactor partners of SPP-like_7942_ based on the Clusters of Orthologous Groups (COGs) classification. Purple indicates proteins identified in one replicate, whereas orange indicates proteins identified in two replicates.

We next used bacterial two-hybrid (B2H) assays to validate a subset of putative interacting proteins, focusing on candidates that have phosphorylated compounds as substrates or products of their reactions (Table S6; Figs. 4 and S9). For each SPP-like_7942_–interactor pair, we tested all possible combinations between these genes fused to T18 or T25 sequences, and at the N-terminus or the C-terminus, since fusion tags could influence protein folding and interaction outcomes. The majority of the hits validated by B2H were proteins that either play a direct role in metabolic functions and/or which themselves are known to be regulated by phosphorylation (Fig. 4). Additionally, KaiB, a core component of cyanobacterial circadian clock (Iwase et al., 2005), and PlmA, a protein involved in the cyanobacterial nitrogen regulatory network (Labella et al., 2016) showed strong interactions with SPP-like_7942_ in B2H assays (Table S7; Fig. S9). Potential substrates of SPP-like_7942_ were inferred based on the substrates and products associated with these putative interactors: Upp - PRPP (phosphoribosyl diphosphate) and UMP (uridine monophosphate); Prs - R5P (ribose-5-phosphate) and PRPP; for SerA - 3PGA (3-phosphoglycerate); and PyrG - uridine triphosphate (UTP) and cytosine triphosphate (CTP) (see Table S7 for more information). Many of these metabolites are involved in nucleotide metabolism, whereas 3PGA is a key intermediate of the Calvin-Benson-Bassham (CBB) cycle.

**Fig. 4.**
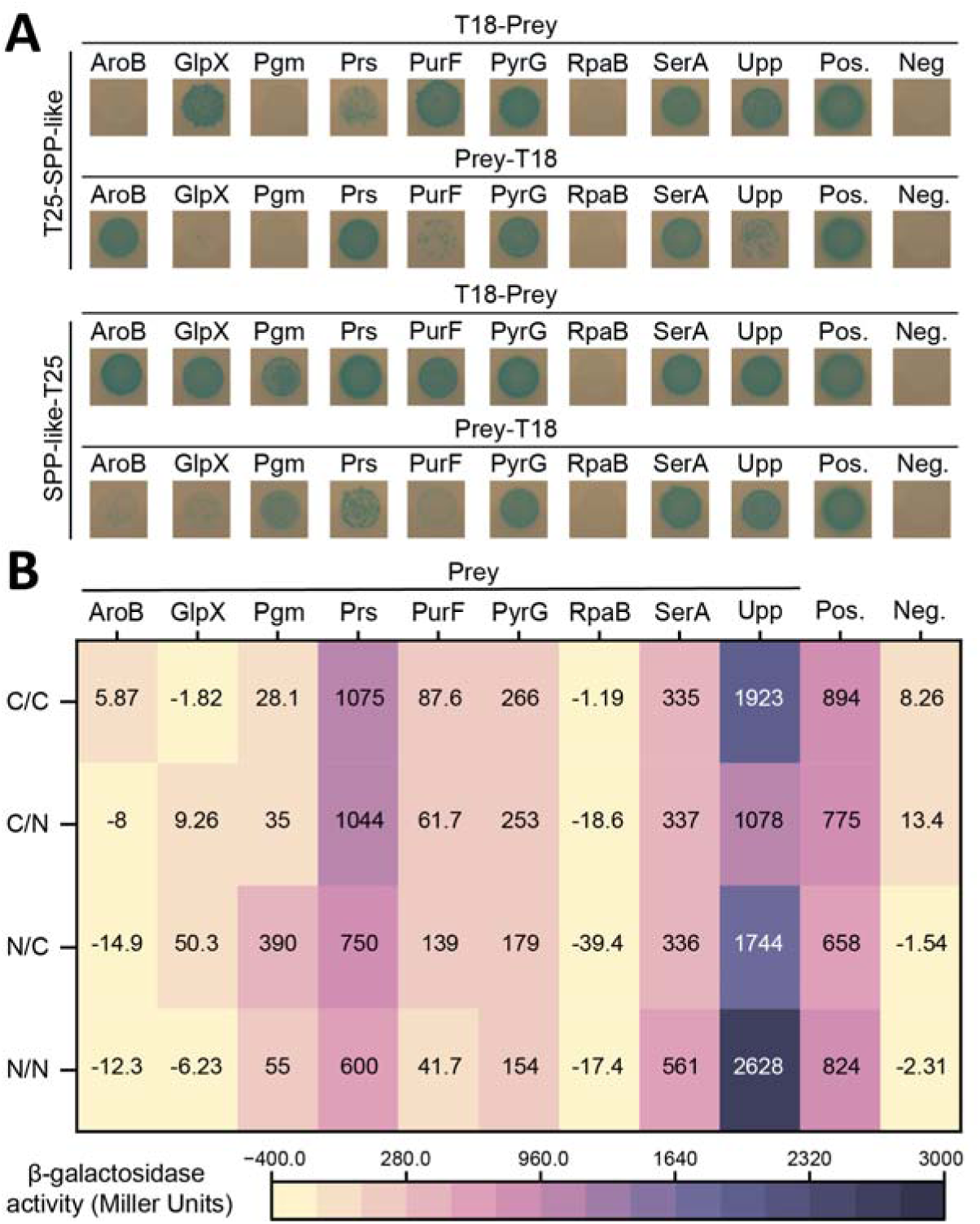
Confirmation of interactions between SPP-like_7942_ and selected candidates from the pull-down by the bacterial two-hybrid system. **(A)** Bacterial two-hybrid analysis of SPP-like_7942_ protein-protein interactions with selected candidates from the heatmap shown in Fig. 3A. *E. coli* strain BTH101 was co-transformed with plasmids encoding the indicated fusions to adenylate cyclase fragments T18 and T25. Colonies were spotted onto selective plates containing IPTG and X-Gal. Blue colonies indicate positive interactions between each pair of fusion proteins. **(B)** Heat map of the quantification of the interaction strength using a β-galactosidase assay. Miller units measured for β-galactosidase activity correlate with the blue colonies observed in the plate assay. Averages of ≥3 independent biological replicates are shown. C/C, T25-SPP-like (bait tagged at the C-terminus) and T18-prey (prey tagged at the C-terminus); C/N, T25-SPP-like (bait tagged at the C-terminus) and prey-T18 (prey tagged at the N-terminus); N/C, SPP-like-T25 (bait tagged at the N-terminus) and T18-prey (prey tagged at the C-terminus); N/N, SPP-like-T25 (bait tagged at the N-terminus) and prey-T18 (prey tagged at the N-terminus); Pos., positive (leucine zipper motif); Neg., negative (linker motif). The β-galactosidase activity values in panel B correspond to the interactions shown in panel A.

### 2.5. SPP-like_7942_ dephosphorylates key phosphorylated metabolites from central carbon metabolism

To identify potential substrates of SPP-like_7942_, we first screened for phosphatase activity against a larger array of putative substrates using excess enzyme. Guided by the list of potential metabolites in Table S6 and the B2H results (Figs. 4 and S9), we tested R5P, PRPP, G3P (glyceraldehyde 3-phosphate), F6P, FBP, G1P, G6P, UTP, CTP, ATP, UMP, and 3PGA as potential substrates for the phosphatase activity of SPP-like_7942_ and SPP_6803_. Some potential phosphorylated substrates (*e.g.*, 3-deoxy-D-arabino-hept-2-ulosonate 7-phosphate) were excluded because they were either commercially unavailable or prohibitively expensive.

SPP-like_7942_ released high amounts of free phosphate from G3P, S6P, and 3PGA, but not from the other metabolites tested (Fig. 5A). In contrast, SPP_6803_ released appreciable phosphate from G6P and S6P, while exhibiting minimal phosphatase activity on R5P, G3P, and ATP, even with large additions of enzyme (30 µg; Fig. 5B). The broad substrate profiles of SPP_6803_ and SPP-like_7942_ align with the well-recognized substrate promiscuity and evolvability of HAD/sugar phosphatases. Enzymes in this family commonly act on diverse phosphorylated sugars and nucleotides, and even small changes in active-site geometry or cap-domain residues can shift specificity between larger hexose phosphates (*e.g.*, G6P, S6P) and smaller triose phosphates (*e.g.*, G3P, 3PGA) (Huang et al., 2015; Kuznetsova et al., 2006; Liu et al., 2025). Interestingly, unlike in the pNPP phosphatase assay, SPP-like_7942_ exhibited higher activity at pH 7.0 than at pH 4.0 for most metabolites tested (Fig. 5A) (Mercan & Bennett, 2010). Thus, the fact that SPP-like_7942_ shows maximal pNPP activity at acidic pH (Fig. 2C) but higher activity toward several physiological substrates at pH 7.0 (Fig. 5A) is most congruent with the fact that enzyme activity on the non-physiological substrate pNPP often diverges from that observed with native metabolites (Mercan & Bennett, 2010). Taken together with its likely cytosolic localization (Figs. 3B and S8), SPP-like_7942_ is most likely physiologically active under neutral pH, despite the tolerance of this protein to denaturation under acidic conditions (Fig. 2E).

**Fig. 5.**
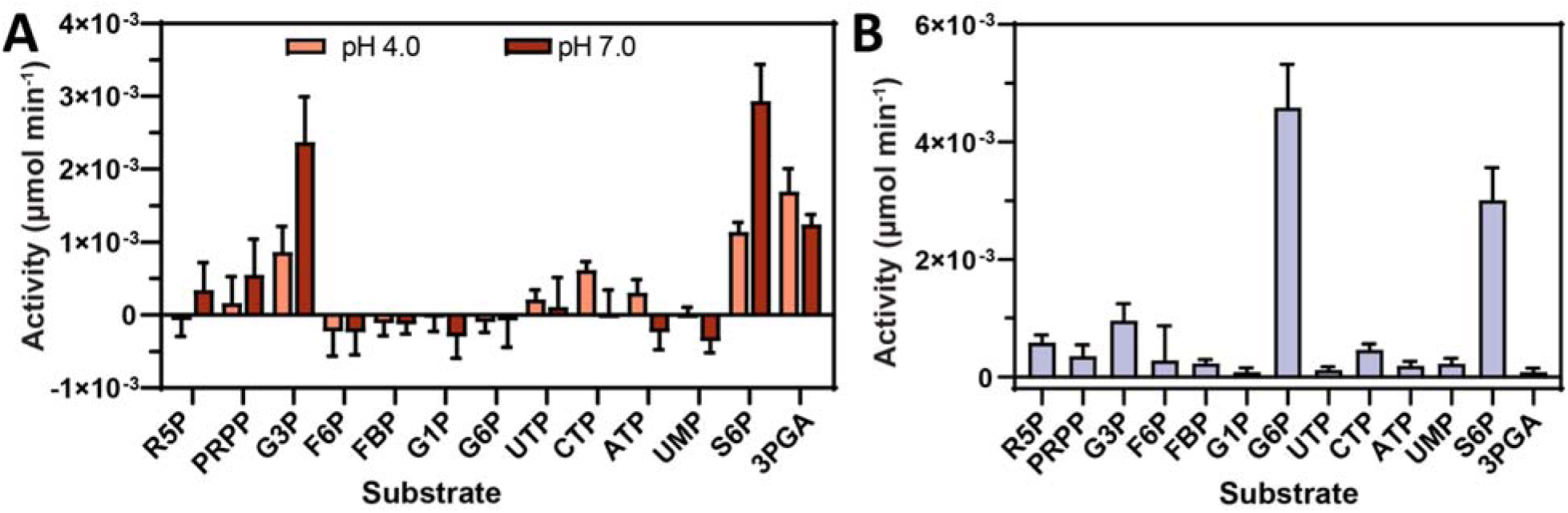
Screening for potential substrates of SPP-like_7942_ and SPP_6803_. **(A)** Phosphatase activity of purified SPP-like_7942_ on various phosphorylated substrates under physiological conditions (pH 7.0 and RT) and under acidic conditions (pH 4.0 and RT). **(B)** Phosphatase activity of purified SPP_6803_ on various phosphorylated substrates under physiological conditions (pH 7.0 and RT). Enzyme activities were calculated as the µmol of inorganic phosphate (as estimated using a commercial kit) released per min in a 100 µL reaction containing 5 mM MgCl_2_, 50 mM HEPES (pH 7.0), 500 µM of each substrate, and 30 µg of enzyme. **(A, B)** Averages of ≥3 independent biological replicates are shown + SD. R5P, ribose 5-phosphate; PRPP, phosphoribosyl pyrophosphate; G3P, glyceraldehyde 3-phosphate; F6P, fructose 6-phosphate; FBP, fructose 1,6-bisphosphate; G1P, glucose 1-phosphate; G6P, glucose 6-phosphate; UTP, uridine triphosphate; CTP, cytidine triphosphate; ATP, adenosine triphosphate; UMP, uridine monophosphate; S6P, sucrose 6-phosphate; 3PGA, 3-phosphoglycerate.

Preliminary data suggested that SPP-like_7942_ may act on substrates beyond small-molecule metabolites, consistent with reports that some HAD phosphatases can moonlight as protein phosphatases (Gohla, 2019). During phosphatase assays using the PiPer™ Phosphate Assay Kit, SPP-like_7942_ displayed a strong background signal in the absence of added substrate (Fig. S10A). In the presence of PiPer™ enzymes, the background signal of free phosphate increased in proportion to the amount of purified SPP-like_7942_ added to the reaction, whereas background phosphate levels were constant regardless of the amount of SPP_6803_ enzyme added (Fig. S10A). Similarly, heat inactivation of the phosphatase reaction prior to phosphate quantification reduced the SPP-like_7942_ background signal several-fold but had no effect on SPP_6803_ (Fig. S10B), indicating that the signal likely arises from intrinsic activity of SPP-like_7942_ rather than co-purified inorganic phosphate. Specifically, the maltose phosphorylase (MP) enzyme provided with the PiPer™ kit was strongly correlated with the release of phosphate when co-incubated with SPP-like_7942_ (Fig. S10C). This suggests that SPP-like_7942_ may have protein phosphatase activity, which is interesting in the context of the several metabolic enzymes and transcription factors that were identified in the SPP-like interactome and validated by B2H (Figs. 3 and 4). However, SPP-like_7942_ protein phosphatase activity remains speculative at this time and would require validation with a verified interaction partner native to *S. elongatus* PCC 7942.

### 2.6. The substrate range for SPP-like_7942_ is broader than S6P

To gain insight into the physiological relevance of SPP-like_7942_ activity in cyanobacteria, we determined substrate kinetics for the metabolites identified as potential substrates (Fig. 6). Comparing enzyme kinetic parameters such as *K*_m_ and *k*_cat_ with intracellular metabolite concentrations is a widely used strategy to assess *in vivo* catalytic potential, as enzyme activity can only be physiologically relevant when substrate levels approach or exceed their *K*_m_.

**Fig. 6.**
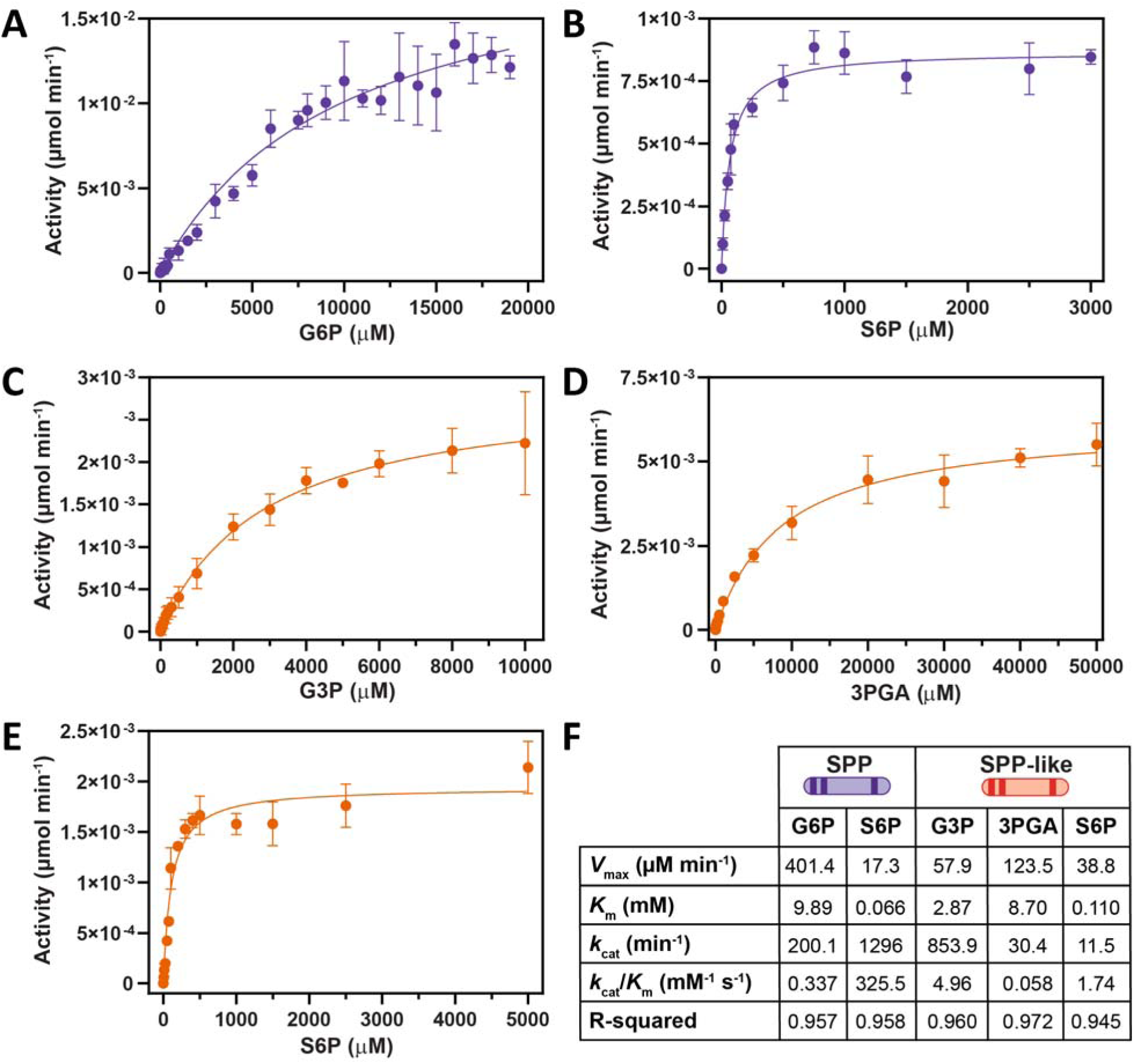
Kinetic parameters of SPP_6803_ and SPP-like_7942_ with different substrates. Michaelis-Menten kinetics of SPP_6803_ with **(A)** G6P, and **(B)** S6P as substrates, respectively. Enzyme activity was calculated as µmol of inorganic phosphate released per min in a 100 µL reaction containing 5 mM MgCl_2_, 50 mM HEPES (pH 7.0) and SPP_6803_ (3 µg for G6P and 0.02 µg for S6P). Michaelis-Menten kinetics of SPP-like_7942_ with **(C)** G3P, **(D)** 3PGA and **(E)** S6P as substrates, respectively. Enzyme activity is calculated as µmol of inorganic phosphate released per min in a 100 µL reaction containing 5 mM MgCl_2_, 50 mM HEPES (pH 7.0) and SPP-like_7942_ (0.1 µg for G3P, 6 µg for 3PGA and 5 µg for S6P). **(F)** Steady-state kinetic constants of the reactions catalyzed by SPP_6803_ and SPP-like_7942_ with various phosphorylated substrates shown in panels A-E performed in 10 mM sodium HEPES buffer (pH 7.0) containing 5 mM MgCl_2_. **(A-E)** Averages of ≥3 independent biological replicates are shown ± SD.

S6P emerged as the substrate with the highest apparent affinity for both enzymes. As expected, SPP_6803_ exhibited robust activity on S6P, with a calculated *K*_m_ of 0.066 mM (Figs. 6B and 6F), broadly consistent with the high substrate specificity previously reported for SPP_6803_ (*K*_m_ = 7.5 µM) (Lunn, 2002). In addition to its known activity toward S6P, SPP_6803_ also dephosphorylated G6P with an apparent *K*_m_ of 9.89 mM (Fig. 6A). This *K*_m_ is substantially higher than estimated intracellular G6P concentrations in both *S. elongatus* PCC 7942 (3.63 mM at high CO□ and 4.05 mM at low CO□) and *Synechocystis* sp. PCC 6803 (0.23 mM at high CO□ and 0.24 mM at low CO□) (Jablonsky et al., 2016; Jablonsky et al., 2014). In both organisms and under both growth conditions, intracellular G6P levels are well below the *K*_m_ of SPP_6803_ (Figs. 6A and 6F), suggesting that SPP_6803_-mediated G6P hydrolysis may be a minor or insignificant reaction *in vivo*. That said, G6P transient accumulation is observed during rapid metabolic reprogramming such as dark-to-light transitions (Shinde et al., 2020; Tanaka et al., 2023), or under nitrogen starvation conditions (Doello et al., 2025). Under these scenarios, temporary increases in G6P availability may partially overcome the high *K*_m_ barrier, which could permit measurable catalytic activity of SPP_6803_ against G6P.

Despite the mutations in residues mapped to the sucrose-binding site and the low sequence similarity (Figs. 1C and S3), SPP-like_7942_ exhibited a similar *K*_m_ of 0.11 mM for S6P (Figs. 6E and 6F), the highest affinity among the metabolites we tested. Notably, the *K*_m_ of SPP-like_7942_ for S6P is generally lower than those reported for SPP homologs from *Arabidopsis* (*K*_m_ = 0.73-3.46 mM) (Albi et al., 2016), from sugarcane and carrot (*K*_m_ = 0.13-0.17) (Hawker & Hatch, 1966), and potato (*K*_m_ = 0.148 mM) (Chen, 2005), but higher than the *K*_m_ reported for rice (*K*_m_ = 0.065 mM) (Echeverría & Salerno, 1994). Although intracellular S6P concentrations are not published in cyanobacteria, the similar *K*_m_ of SPP-like_7942_ to SPP_6803_ and higher apparent affinity of SPP-like_7942_ in comparison to plants suggest that S6P is likely a target of SPP-like_7942_ under physiological conditions. SPP-like_7942_ exhibits a lower *k*_cat_ toward S6P relative to the canonical SPP_6803_, suggesting that SPP-like_7942_-mediated S6P dephosphorylation in *S. elongatus* PCC 7942 may be most relevant when SPP expression from the SPPb is low or absent, such as under non-salt-stress conditions (Liang et al., 2020). In this context, SPP-like_7942_ may function as a low-capacity, auxiliary enzyme that sustains sucrose synthesis without driving high flux.

If the dominant function of SPP-like proteins is solely to perform the metabolic function of dephosphorylation of S6P, this would raise questions about their evolutionary relationships. For example, why would cyanobacterial genomes frequently encode both canonical SPP genes and SPP-like genes (Figs. 1B and S2)? This would suggest evolutionary conservation of redundant functions, and would be particularly unexpected if, as with SPP-like_7942_, the SPP-like family is generally a similar affinity and lower flux enzyme class. Indeed, it is notable that SPP-like homologs are more widespread than canonical SPP genes, and that many cyanobacterial genomes encode one or more SPP-like proteins while lacking SPPu or SPPb altogether (Figs. 1B, 1C, S2 and Table S1). Furthermore, SPP genes often exhibit stress-responsive expression patterns, such as under hyperosmotic conditions (Klahn et al., 2021; Liang et al., 2020), while this is not generally true of SPP-like expression patterns (Billis et al., 2014). Taken together, one interpretation is that SPP-like proteins may serve physiological roles that are distinct from the well-characterized function of SPP in sucrose metabolism.

Consistent with other possible cellular functions, we observed phosphatase activity of SPP-like_7942_ toward two key metabolites from central carbon metabolism (G3P and 3PGA) in addition to the putative protein phosphatase activity described in Section 3.5. SPP-like_7942_ displayed a high catalytic efficiency (*k*_cat_/*K*_m_) with G3P (Figs. 6C and 6F), yet the intracellular concentration of G3P in *S. elongatus* PCC 7942 is estimated at only ∼51.3 µM (Gao et al., 2016), which is substantially below the measured *K*_m_ of 2.87 mM. Furthermore, accumulation of G3P to levels approaching the *K*_m_ seems unlikely *in vivo* because of its aldehyde toxicity (Xiong et al., 2015).

By contrast, the apparent *K*_m_ of SPP-like_7942_ against 3PGA (8.7 mM; Fig. 6D) overlaps with reported intracellular concentrations. Intracellular 3PGA concentrations in *S. elongatus* PCC 7942 range from 4.99 mM at high CO to 13 mM at low CO (Jablonsky et al., 2014), with similar levels in *Synechocystis* sp. PCC 6803 (Jablonsky et al., 2016). Similarly, an intracellular concentration of ∼12 mM 3PGA has been reported in *Synechocystis* sp. PCC 6803 in the dark (Tanaka et al., 2023), and increased levels have also been observed during nitrogen limitation (Lucius & Hagemann, 2024). These reports further support the possibility that SPP-like_7942_-mediated turnover of 3PGA could become physiologically significant under stress or metabolic imbalance.

The broad distribution and evolutionary conservation of SPP-like family members suggest that they might play distinct cellular roles compared to the better-characterized SPP enzymes. Nominally, a simple interpretation is that SPP-like enzymes may maintain basal flux through the sucrose biosynthesis pathway under non-stress conditions where SPP levels are low in the cell. Alternatively, given the multiple regulatory roles of 3PGA in central carbon metabolism, including Rubisco regulation (Lobo et al., 2024), activation of ADP-glucose pyrophosphorylase (GlgC), and glycogen synthesis (Lee et al., 2025), it is feasible that SPP-like_7942_ may influence carbon partitioning during carbon limitation or other stress conditions where 3PGA levels spike. However, our preliminary analysis of photosynthetic activity in Δ*spp-like*_7942_ strains under a few laboratory growth conditions did not reveal obvious fitness defects in photosynthetic performance or cellular growth rates under laboratory conditions (Figs. S11 and S12). The lack of a clear photosynthetic phenotype in Δ*spp-like*_7942_ strains would suggest that either SPP is able to compensate for its loss, or that SPP-like_7942_ exerts a greater influence on carbon metabolism under environmental conditions that were not tested in this study. A more speculative possibility is that SPP-like_7942_ exhibits physiologically relevant phosphatase activity against one or more proteins in *S. elongatus* PCC 7942 (Fig. S10).

## 3. Conclusion

Our findings suggest that the cyanobacterial SPP-like family of HAD subfamily IIB proteins is widespread and is both functionally and genetically distinct from the better-characterized SPP family. Despite divergence in key active site residues, SPP-like_7942_ retains bona fide phosphatase activity toward S6P, indicating other SPP-like family members may also target S6P. In addition to phosphatase activity on S6P, SPP-like_7942_ also exhibits activity toward 3PGA within a physiologically relevant concentration range, suggesting potential implications for regulation of the CBB cycle. One interpretation of our data is that the divergence of SPP-like from canonical SPPs may represent an evolutionary trade-off between S6P specificity and a wider pool of substrate targets. The putative protein phosphatase activity of SPP-like_7942_, together with its interactions with several transcriptional regulators and metabolic enzymes, further suggests a broader regulatory role in cyanobacteria. The conservation of SPP-like proteins across diverse cyanobacterial species and ecosystems strongly supports the idea that their functions are distinct from and non-redundant with those of the canonical SPP proteins. However, further characterization of the Δ*spp-like* mutant under a broader range of environmental conditions will be required to define the physiological role of this protein *in vivo*.

## 4. Data availability

Data will be made available on request. E-supplementary data for this work can be found in e-version of this paper online.

## 5. Acknowledgements

This work was primarily supported by the U.S. Department of Energy, Office of Science, Office of Basic Energy Sciences, United States Department of Energy under Award Number DE-FG02-91ER20021 to D.C.D. Additional support for this work was provided by the National Science Foundation Office of Emerging Frontiers in Research and Innovation, Award #2029374. We would like to thank Dr. Douglas Whitten for critical feedback on experimental setup, mass spectrometry sample processing and guidance with the respective data analysis.

