## Supplementary material for "A novel class of conserved sucrose-phosphate phosphatases highlights the diversity of cyanobacterial sucrose metabolism"

1. **Supplementary texts**

**Text S1 – Blue-native gel electrophoresis**

For native-PAGE, 5 µg of protein samples were mixed to a final volume of 10 µL with NativePAGE™ Sample Buffer (1X), NativePAGE™ 0.002% G-250 Coomassie Sample Additive, and deionized water then loaded onto a NativePAGE™ 4-16% Bis-Tris Gel (BN1002BOX). The NativeMark™ Unstained Protein Standard was used as a molecular weight marker. Electrophoresis was performed at room temperature (RT) with pre-chilled running buffers for 2 h at 150 V. The gel was then subjected to Coomassie Stain (10% v/v acetic acid, 45% ethanol, Coomassie Brilliant Blue 2 g R-250, 0.5 g G-250 per liter) for 30 min with subsequent rounds of destaining to remove background stain (10% v/v acetic acid, 30% v/v ethanol). The destained gel was imaged on a Bio-Rad GelDoc Go Imaging System.

**Text S2 - Structural prediction using AlphaFold2 Multimer**

To predict the oligomeric structures of SPP-like_7942_ and SPP_6803_ AlphaFold2-Multimer was used (Lin et al., 2025; Mirdita et al., 2022). Structures containing 2, 3, 4, and 6 copies of each protein were predicted with 3 recycles with 200 subsequent relaxation iterations. The interface predicted TM (ipTM) score was obtained for each oligomeric state and each protein. Values of ipTM higher than 0.8 indicate high-confidence and high-quality predictions, whereas values lower than 0.6 indicate a likely failed prediction. The top-ranked structure of the SPP-like_7942_ dimer was rendered in Fig. S4A using ChimeraX (Meng et al., 2023).

**Text S3 – Determination of protein phosphatase activity**

To determine whether SPP-like_7942_ exhibits protein phosphatase activity, 1 µL of the three enzymes from the PiPer™ kit (MP, maltose phosphorylase; GO, glucose oxidase; or HRP, horseradish peroxidase) were individually used as substrate in a 100 µL phosphatase assay containing 50 mM HEPES (pH 7.0), 10 mM MgCl₂, and 30 µg of SPP-like_7942_. Reactions were incubated at RT for 10 min and subsequently heat-inactivated at 80 °C for 20 min. 50 µL of inactivated reaction mixture was combined with 50 µL of PiPer™ reagent and incubated at RT for 30 min before fluorescence was measured as described above.

**Text S4 – In-gel protein digestion and extraction**

Briefly, gel bands were dehydrated using 100% acetonitrile and incubated with 10 mM dithiothreitol in 100 mM ammonium bicarbonate, pH ~8.0, at 56 °C for 45 min, dehydrated again and incubated in the dark with 50 mM iodoacetamide in 100 mM ammonium bicarbonate for 20 min. Gel bands were then washed with ammonium bicarbonate and dehydrated again. Sequencing-grade modified trypsin was prepared to 0.005 µg µL^-1^ in 50 mM ammonium bicarbonate and ~100 µL of this was added to each gel band to completely submerge the gel. Bands were then incubated at 37 °C overnight. Peptides were extracted from the gel by water-bath sonication in a solution of 60% acetonitrile/1% trifluoroacetic acid (v/v) and vacuum dried to ~2 µL.

**Text S5 – LC–MS/MS analysis and LC–MS/MS data processing**

All samples were reconstituted in 20 µL of 2% acetonitrile/0.3% trifluoroacetic acid (v/v) and an injection of 5 µL (10 µL for bead samples) was made using an Easy-nLC 1000 system (Thermo Fisher Scientific, Somerset, NJ) onto an Acclaim PepMap RSLC 0.1 mm x 20 mm C18 trapping column (Thermo Fisher Scientific, Somerset, NJ) and washed for ~5 min with buffer A. Bound peptides were then eluted over 35 min onto an Acclaim PepMap RSLC 0.075 mm x 250 mm resolving column (Thermo Fisher Scientific, Somerset, NJ) with a gradient of 5% buffer B to 38% buffer B over 24 min, ramping to 90% buffer B at 25 min and held at 90% buffer B for the duration of the run (buffer A = 99.9% water/0.1% formic acid (v/v), buffer B = 80% acetonitrile/0.1% formic acid/19.9% water (v/v)) at a constant flow rate of 300 nL min^-1^. Column temperature was maintained at a constant temperature of 50 °C using an integrated column oven (PRSO-V1, Sonation GmbH, Biberach). Eluted peptides were sprayed into a Q-Exactive mass spectrometer (Thermo Fisher Scientific, Somerset, NJ) using a FlexSpray spray ion source. Survey scans were taken in the Orbitrap (70000 resolution, determined at m/z 200), and the top 15 ions in each survey scan selected for higher-energy collision-induced dissociation (HCD), with fragment spectra acquired at 17,500 resolution. The resulting MS/MS spectra were converted to peak lists using Mascot Distiller v2.8.1 and searched against a database containing all *S. elongatus* PCC 7942 protein sequences available from Uniprot appended with common laboratory contaminants (downloaded from www.thegpm.org, cRAP project) using the Mascot searching algorithm v2.8.0.1. The Mascot output was then analyzed using Scaffold v5.1 to probabilistically validate protein identifications. Assignments validated using the Scaffold 1% false discovery rate (FDR) confidence filter are considered true.

MaxQuant/Andromeda (version 2.0.3.1) (Tyanova et al., 2016) was used to process raw files from the Q-Exactive mass spectrometer and search peak lists against a database consisting of Uniprot *S. elongatus* PCCC 7942 proteome (UP000002717, total 2,657 entries, downloaded at 10/15/2024). Enzyme specificity was set as full cleavage by trypsin with two maximum missed cleavage sites permitted. Carbamidomethylation (Cys) was set as fixed modification, whereas dynamic modifications were set as oxidation (Met), deamidation (Asn/Gln), and acetylation (Protein N-terminal). MaxQuant used 4.5 ppm main search tolerance for precursor ions, 20 ppm MS/MS match tolerance, searching top 12 peaks per 100 Da. False discovery rates (FDRs) for both protein and peptide were 0.01 with a minimum of seven amino acid peptide length, and the minimum score for peptides was set to >40. Label-free quantification was enabled with minimum 2 LFQ ratio counts and a fast LFQ option. The mass spectrometry proteomics data has been deposited to the ProteomeXchange Consortium via the PRIDE partner repository with the dataset identifier PXD079470 and 10.6019/PXD079470 (Perez-Riverol et al., 2022).

LFQ values were log_2_ transformed and missing intensity values were imputed using the “min” method, which replaces the missing values by the smallest non-missing value in the data set (Lazar et al., 2016). Only proteins with at least two unique peptides and two spectral counts were considered identified in an individual sample. The log_2_ ratio of the MaxQuant LFQ intensities was used as metrics to determine enrichment in the 6xHisTag-SPP-like samples over a control sample with only beads (log_2_ LFQ SPP-like/control) (Old et al., 2005). Unpaired two-sided Student’s t-tests were performed to determine the statistical significance between 6xHisTag-SPP-like and control samples. Proteins were considered candidate interactors when log2 LFQ SPP-like/control was >1.5 and p-value in an unpaired Student’s t-test was <0.05.

**Text S6 - Strains and culture conditions**

*S. elongatus* PCC 7942 cultures were grown in BG11 medium supplemented with 1 g L^−1^ HEPES to a final pH of 8.3 with NaOH. Flasks were cultured in a Multitron incubator (Infors HT) at 32 °C under ambient air CO_2_ (LC) or supplemented with 2% CO_2_ (HC) with ~150 μmol photons m^−2^ s^−1^ of light (high light – HL) or 17 μmol photons m^−2^ s^−1^ of light (low light – LL) provided by Sylvania 15 W Gro-Lux fluorescent bulbs and shaken at 150 rpm. Cultures were back-diluted daily to an OD_750_ of 0.3 for at least 3 days before experiments or theophylline (theo) induction. Where appropriate, 1 mM theo was used to induce *spp-like* gene expression or 150 mM NaCl was used to induce osmotic stress. Spectinomycin (Sp; 100 μg mL^−1^) was used to maintain WT/Δ*spp-like*, OE-SPP-like-mNG and OE-mNG-SPP-like strains. In all cases, antibiotic selection was removed before performing the experiments to minimize unintended effects.

**Text S7 - Strain construction**

The genomic locus encoding the *spp-like* gene was disrupted by inserting a spectinomycin resistance cassette (*aadA*; Sp^r^). To visualize the subcellular localization of SPP-like_7942_, fluorescent reporter fusions were generated by integrating mNeonGreen (mNG) at either the N-terminus (mNG-SPP-like) or the C-terminus (SPP-like-mNG) of the *spp-like* gene using an NS1 integration plasmid. Plasmid details are provided in Table S3.

**Text S8 - Photosynthetic fluorescence measurements**

Samples containing cyanobacteria (2.5 μg mL^−1^ chlorophyll) were resuspended in fresh medium sparged with 2% CO_2_ in air and dark-adapted for 3 min before measurement (Santos-Merino et al., 2021). The apparent quantum yield of photosystem II (Φ_II_) calculated as (F’_M_ - F_S_)/(F’_M_) was measured using a 1.5 s saturating pulses of actinic light (~5,000 μmol photons m^−2^ s^−1^).

**Text S9 - Microscopy and Image Analysis**

A 2 mL culture aliquot was centrifuged at 10,000 ×g for 5 min, resuspended in 80 μL of BG11 medium, and a 2 μL aliquot was transferred onto a 3% agarose pad. After equilibration for at least 10 min, the pad was placed onto a #1.5 glass coverslip and imaged using a Zeiss Axio Observer D1 inverted microscope equipped with an Axiocam 503 mono camera and a Zeiss Plan-Apochromat 63X/1.4 NA oil-immersion objective.

2. **Supplementary figures**

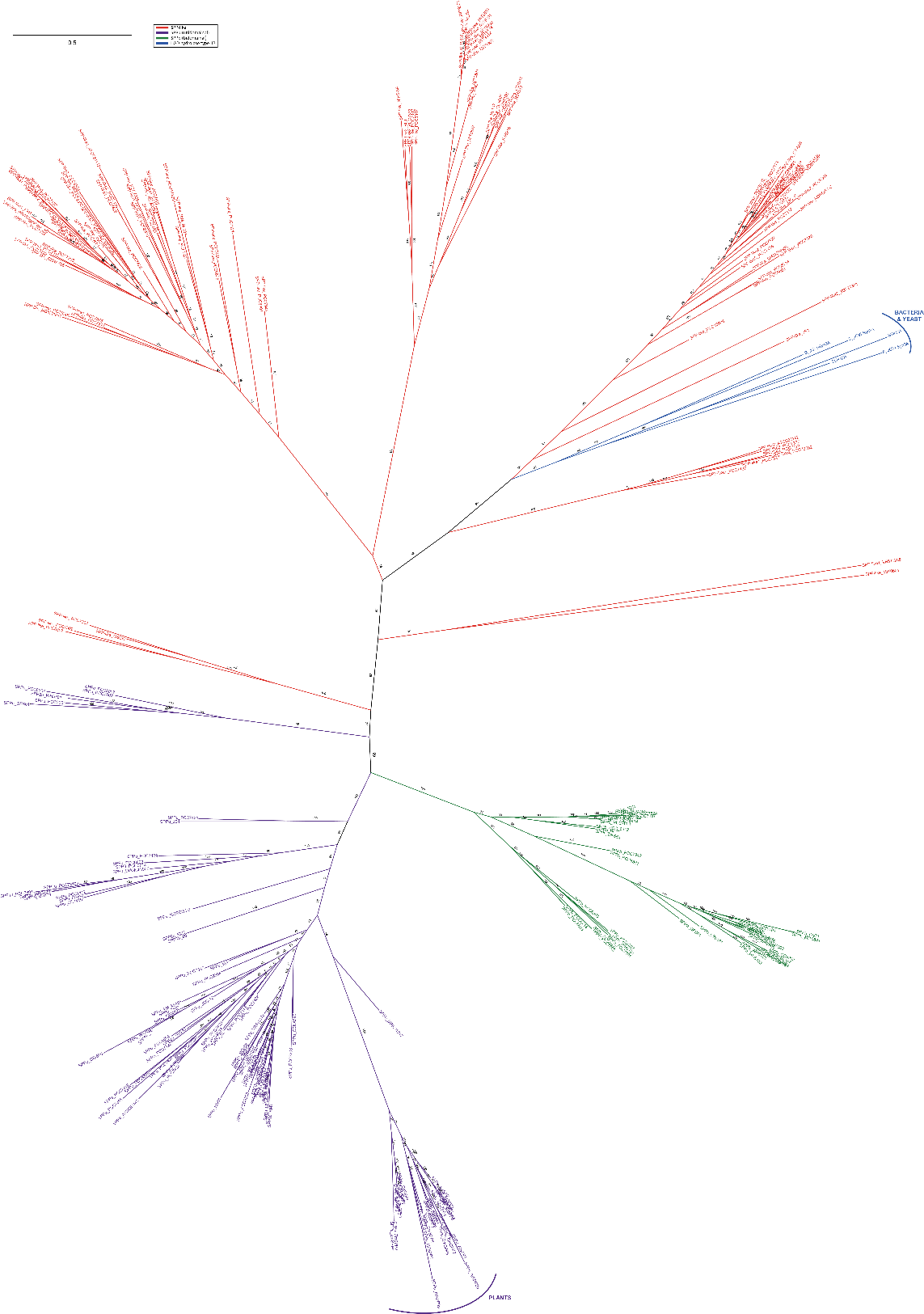

**Figure S1. Diversity of SPP and SPP-like proteins across different groups.** Phylogenetic tree of the evolutionary relationships between SPP unidomainal (SPPu), SPP bidomainal (SSPb) and SPP-like of cyanobacteria and plants, as well as HAD hydrolase type IIB from bacteria and yeast. Numbers at each branch point indicate the bootstrap support values (percentages) calculated from 1,000 replicate trees using IQ-TREE (Trifinopoulos et al., 2016).

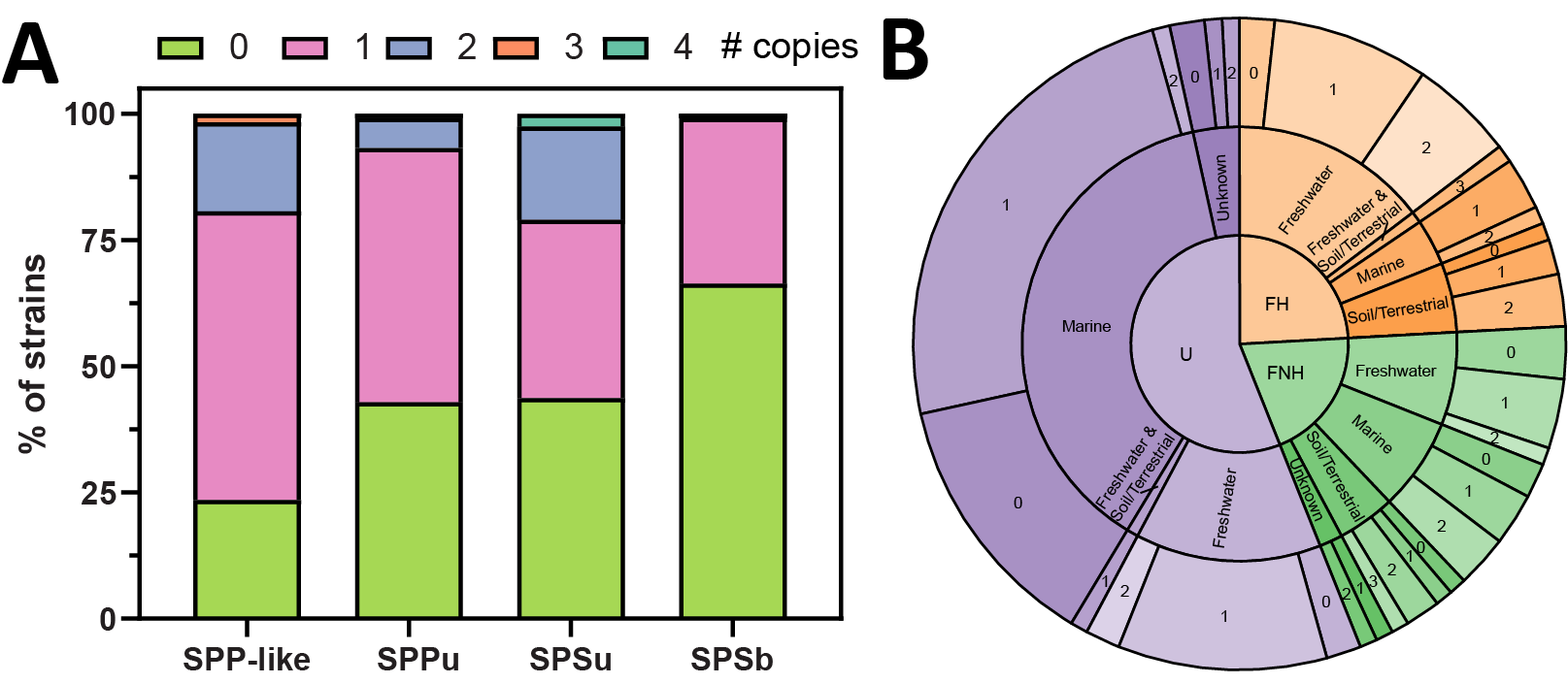

**Figure S2. Distribution of the number of genes encoding SPP-like proteins and other proteins involved in sucrose synthesis. (A)** Bar graph representing the percentage of cyanobacterial genomes listed in Table S1 that contain genes encoding SPP-like, SPPu, SPSu and SPPb. **(B)** Distribution of cyanobacterial genomes containing genes encoding SPP-like according to morphology and habitat. The number of filamentous cyanobacteria is underrepresented in this analysis because sequenced filamentous cyanobacterial genomes are underrepresented in databases (Cornet et al., 2018). U, unicellular; FH, filamentous heterocyst-forming; FNH, filamentous non-heterocyst forming. Of the 119 genomes analyzed, five species lacked any recognizable homolog of SPPu, SPPb, or SPP-like, which may indicate alternative pathways for sucrose metabolism or incomplete/fragmented genomic data (Teikari et al., 2022).

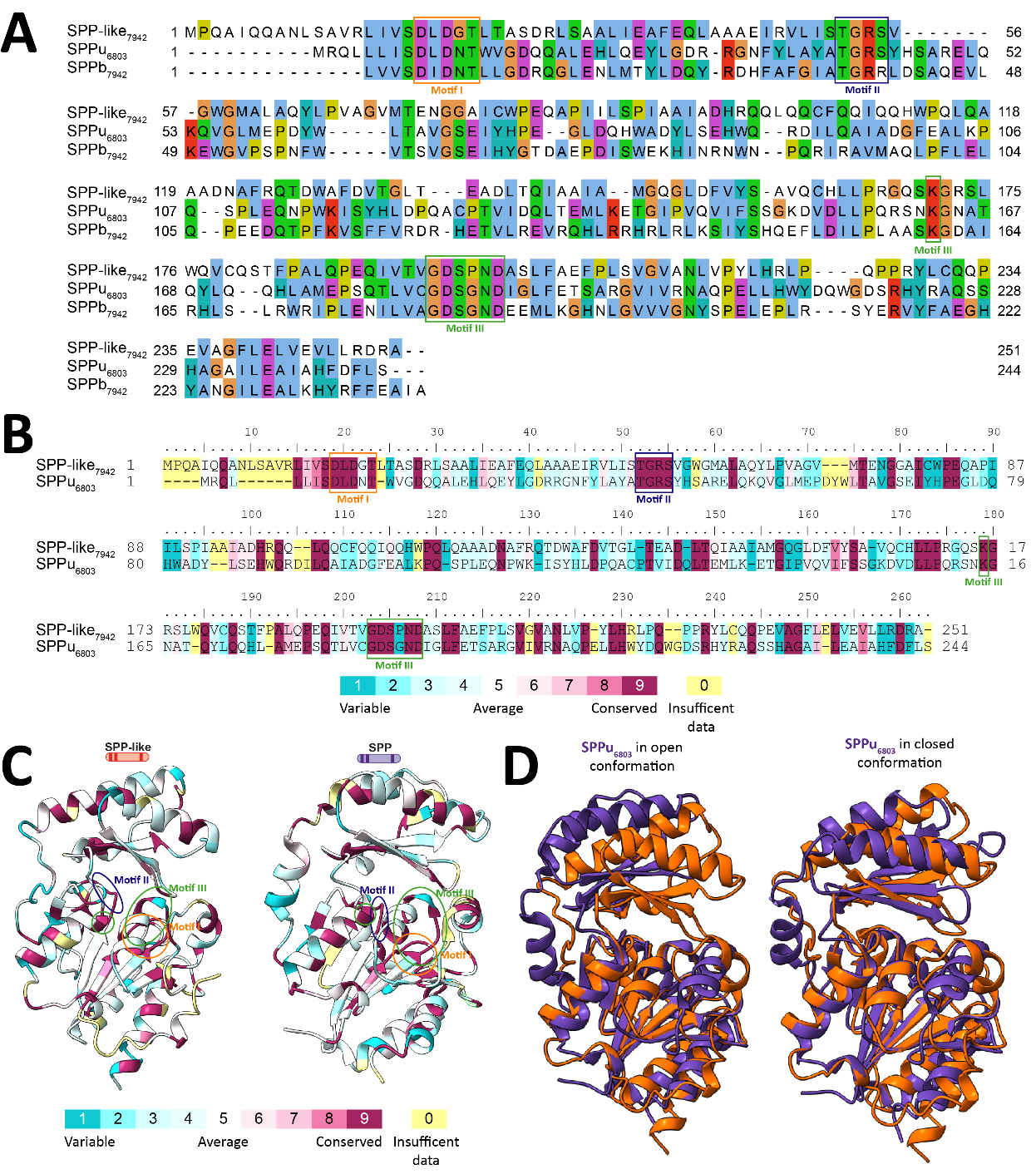

**Figure S3. Low sequence similarity but high structural similarity between SPP-like**_7942_ **and SPP_6803_. (A)** ClustalW alignment of the SPP-like_7942_, SPPu_6803_ and SPPb_7942_ sequences showing conservation of the HAD motifs but variability across the remainder of the sequences**. (B)** Evolutionary conservation of amino acid positions in SPP-like_7942_ and SPPu_6803_, as determined by the ConSurf server. Different colors indicate the variability range of each amino acid position. **(C)** The mutational variability color scale determined by the ConSurf server was mapped onto the protein structures of SPP-like_7942_ and SPPu_6803_. **(D)** Structural comparison of SPP-like_7942_ (orange) and SPP_6803_ (purple), in open conformation (left; PDB ID: 1S2O) and closed conformation (right; PDB ID: 1TJ3) based on previously published structures (Fieulaine et al., 2005).

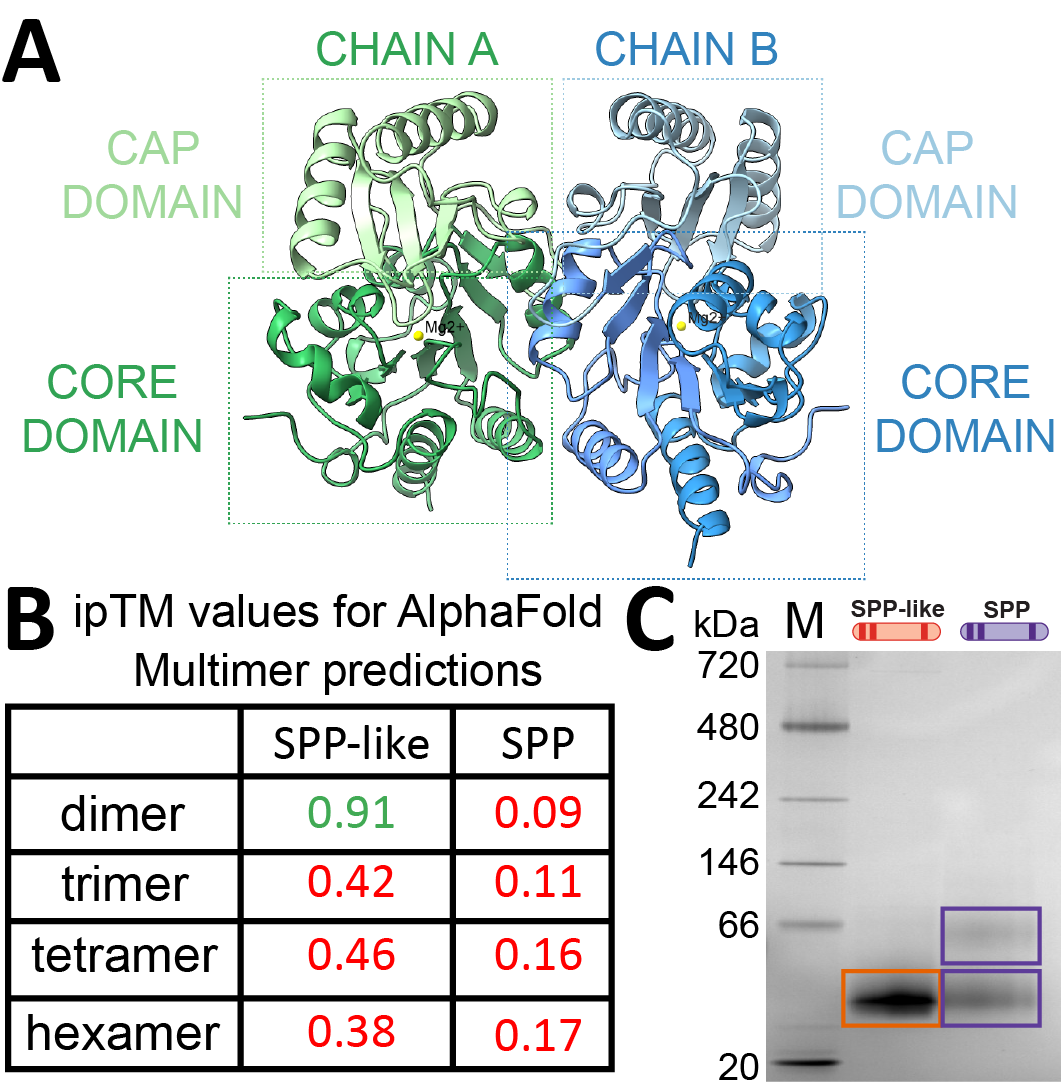

**Figure S4. SPP-like**_7942_ **forms a dimer. (A)** AlphaFold predicted dimeric structure of SPP-like_7942_. **(B)** The predicted dimeric conformation of SPP-like_7942_ has an ipTM value of 0.91, whereas SPP_6803_ has an ipTM value of 0.09. **(C)** Blue-native PAGE analysis of SPP and SPP-like_7942_.

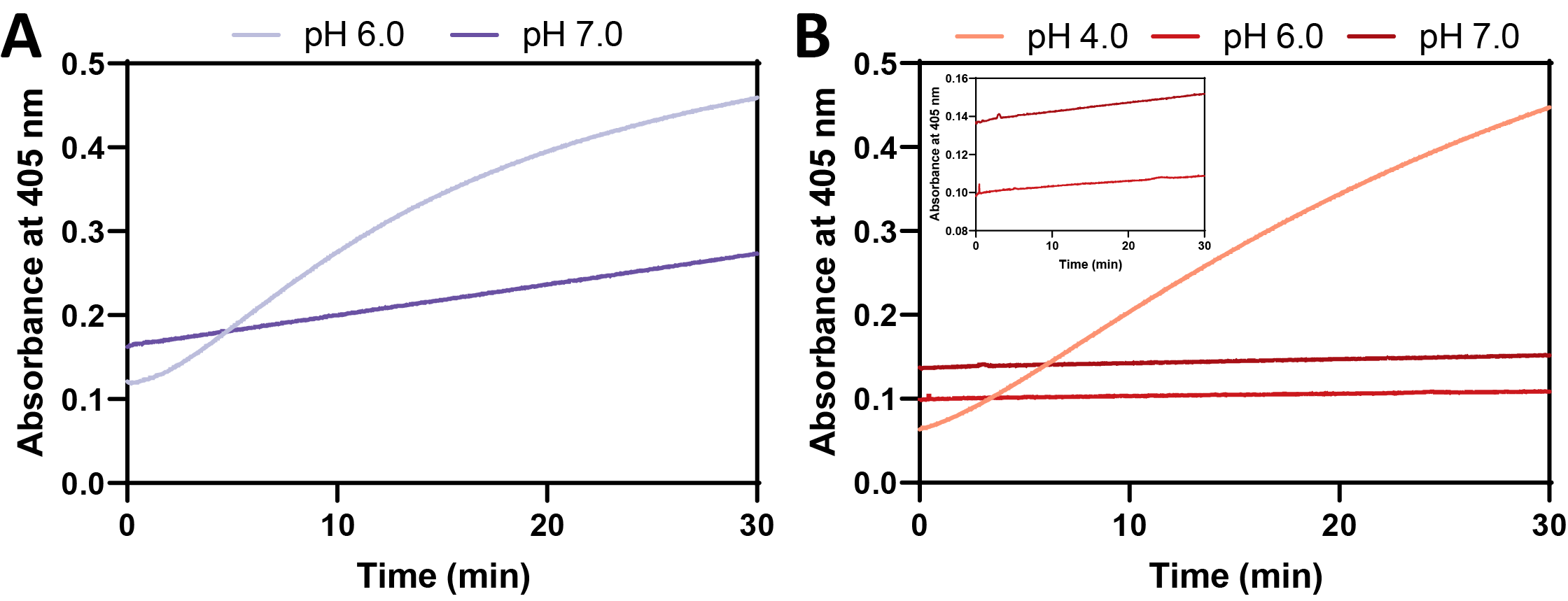

**Figure S5.** **SPP-like exhibits prolonged activity (30 min) across all tested pH conditions.** pNPP phosphatase activity of **(A)** SPP_6803_ and **(B)** SPP-like_7942_ measured by monitoring the formation of pNP at 405 nm under different pH conditions.

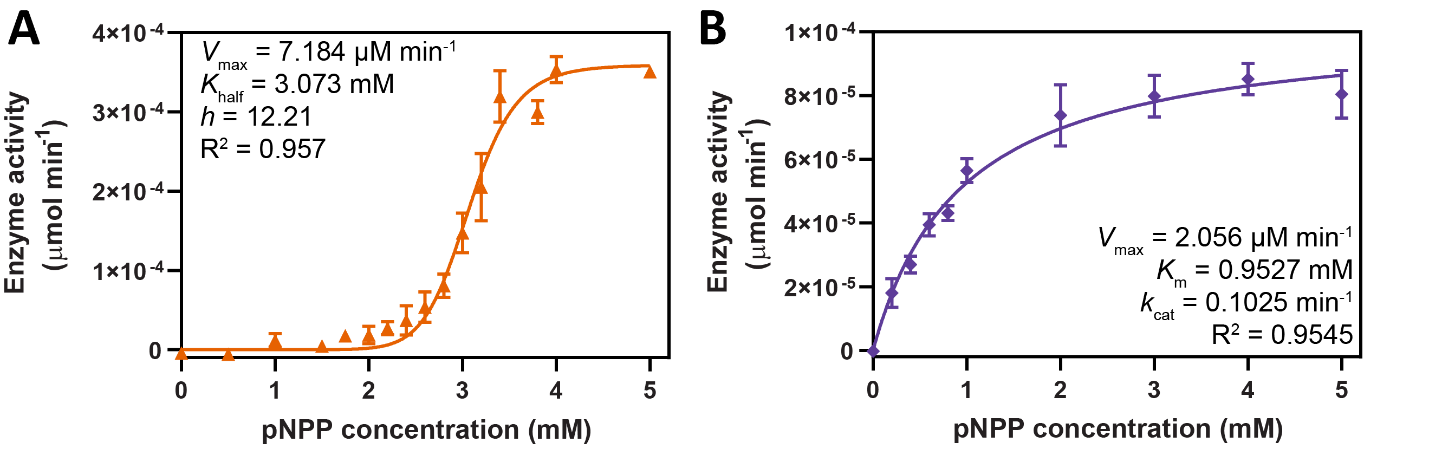

**Figure S6. SPP-like and SPP**_6803_ **enzyme kinetics using the universal substrate pNPP. (A)** Michaelis–Menten kinetics of SPP-like_7942_ using pNPP as a substrate at pH 4.0 and at RT. **(B)** Michaelis–Menten kinetics of SPP_6803_ using pNPP as a substrate at pH 7.0 and at RT. **(A, B)** Averages of ≥3 independent biological replicates are shown ± SD.

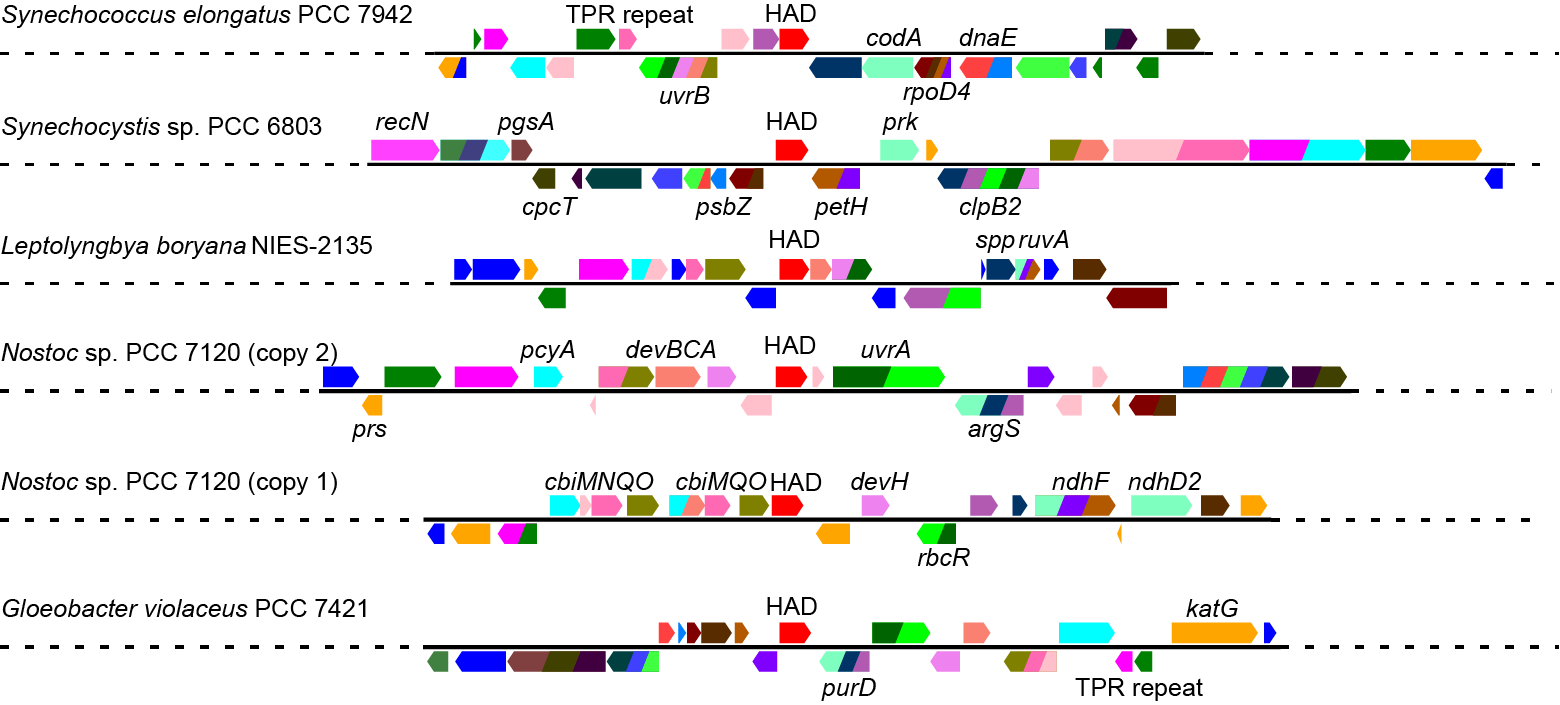

**Figure S7. Gene neighborhood analysis of selected SPP-like sequences from different cyanobacterial species.** The selected SPP-like proteins were chosen from different branches of the phylogenetic trees shown in Figs. 1A and S1. Proteins with unknown functions are not labeled.

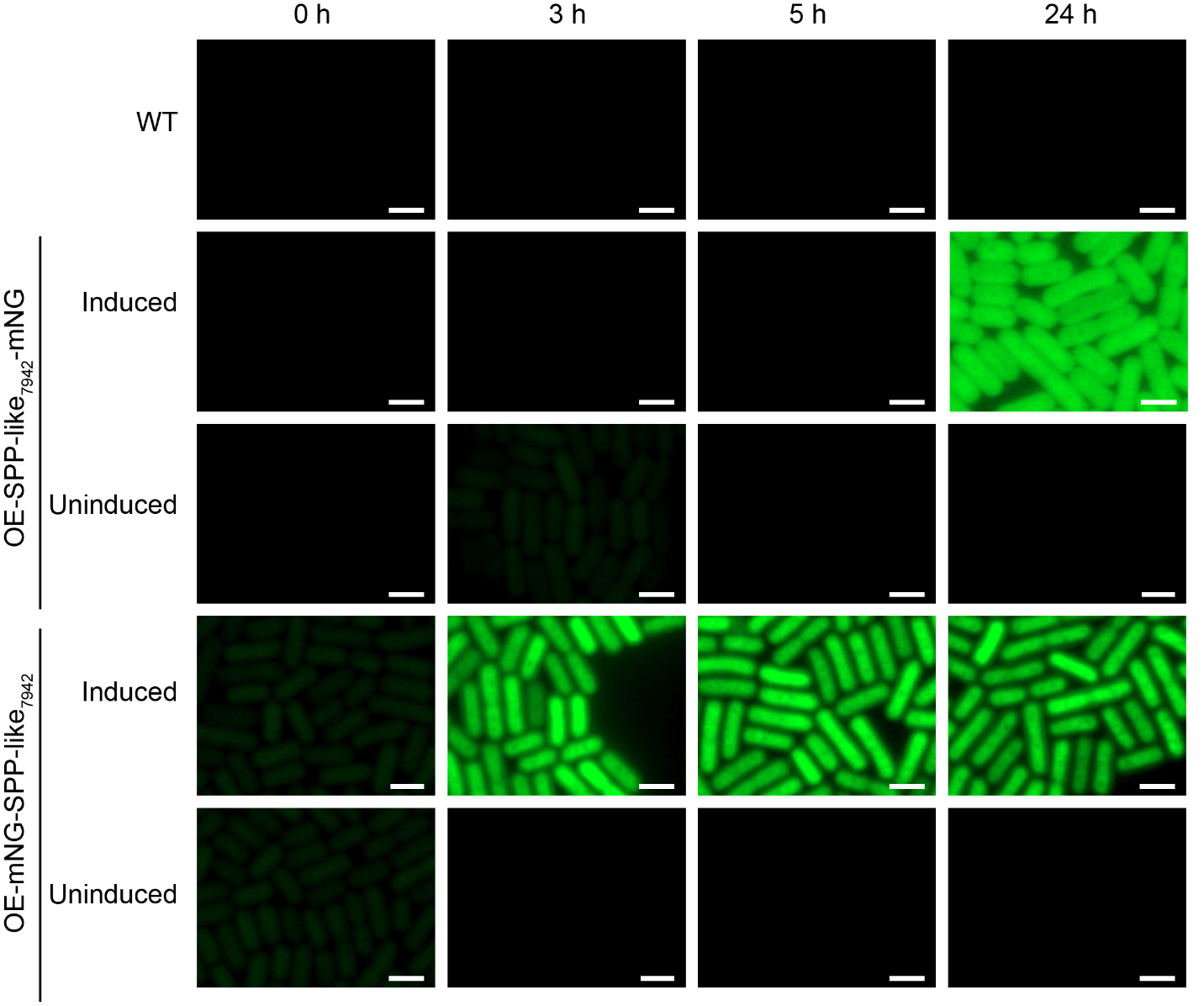

**Figure S8. *In vivo* localization of SPP-like_7942_ using mNeonGreen (mNG).** An additional copy of *spp-like* tagged with mNG at either the C-terminus (OE-SPP-like_7942_-mNG) or the N-terminus (OE-mNG-SPP-like_7942_) was integrated into neutral site 1. Protein expression was controlled using an inducible promoter (P*trc* with a theophylline-dependent riboswitch) by the addition of 1 mM theo. Scale bar: 2 μm.

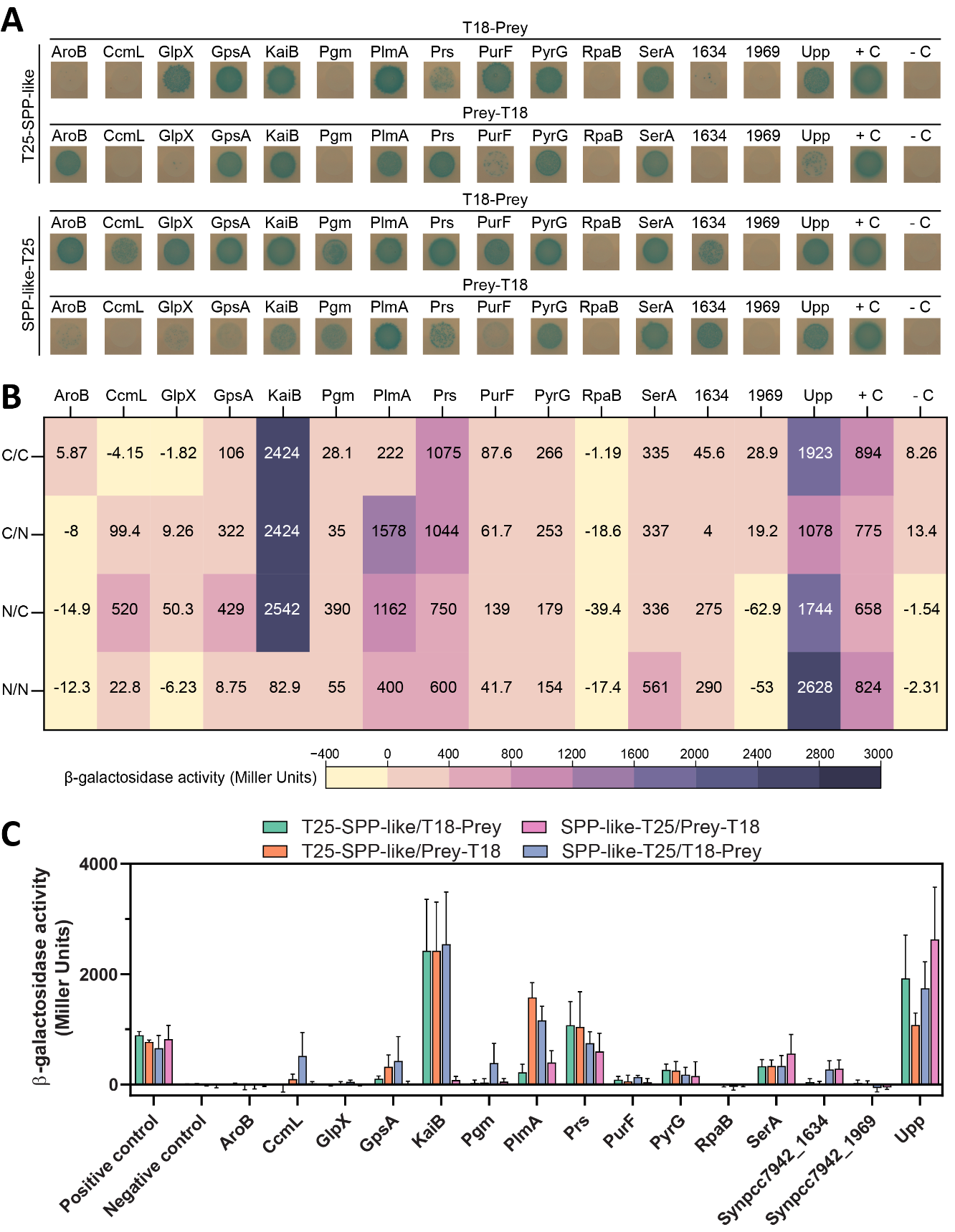

**Figure S9. Confirmation of the interaction between SPP-like_7942_ and selected candidates from the pull-down assay using the bacterial two-hybrid system. (A)** Bacterial two-hybrid analysis of SPP-like_7942_ protein-protein interactions with selected candidates of the heatmap shown in Fig. 4A, and the expanded candidates shown in Fig. 5. *E. coli* strain BTH101 was co-transformed with plasmids encoding the indicated fusions to the adenylate cyclase fragments T18 and T25. Colonies were spotted onto selective plates containing IPTG and X-Gal. Blue colonies indicate positive a positive interaction between each pair of fusion proteins. **(B)** Heat map showing the quantification of interaction strength using a β-galactosidase assay. Miller units measured for β-galactosidase activity correlate with blue colonies observed in plate assay. Averages of ≥3 independent biological replicates are shown. C/C, T25-SPP-like and T18-prey; C/N, T25-SPP-like and prey-T18; N/C, SPP-like-T25 and T18-prey; N/N, SPP-like-T25 and prey-T18. **(C)** Bar graph showing the quantification of interaction strength using a β-galactosidase assay. Miller units measured for β-galactosidase activity correlate with blue colonies observed in plate assay. Averages of ≥3 independent biological replicates are shown + SD.

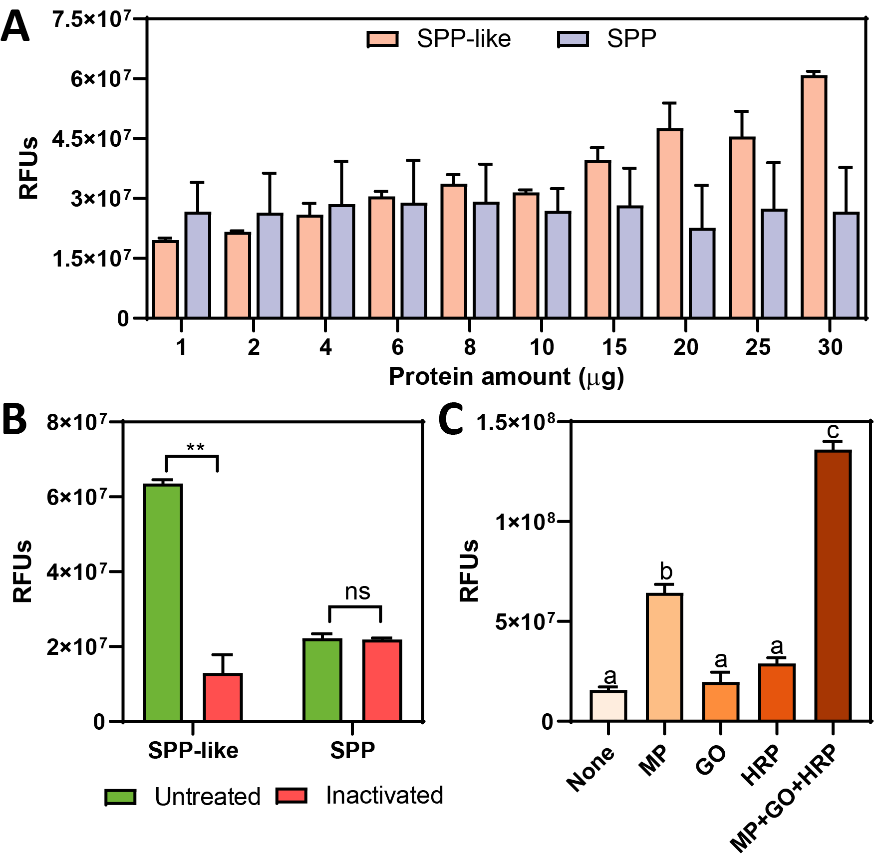

**Figure S10. Background fluorescence signal in phosphatase assays with SPP-like_7942_ and SPP_6803_.** **(A)** Background fluorescence signal in phosphatase assays containing only enzymes (SPP-like_7942_ and SPP_6803_) without any substrate. Averages of ≥3 independent biological replicates are shown + SD. **(B)** Background fluorescence signal in phosphatase assays containing only enzymes (SPP-like_7942_ and SPP_6803_) using either untreated samples or heat-inactivated proteins (20 min at 80 °C). Averages of ≥3 independent biological replicates are shown + SD. Significance was calculated by unpaired Student’s t test relative to uninduced strain as the control. **P < 0.01; ns, not significant. **(C)** Background fluorescence signal in phosphatase assays containing SPP-like_7942_ and the three protein components of the PiPer™ kit (MP, maltose phosphorylase; HRP, horseradish peroxidase; GO, and glucose oxidase), alone or in combination. Averages of ≥3 independent biological replicates are shown + SD. Significance was calculated by one-way ANOVA followed by Tukey’s multiple comparison test. Bars labeled with different letters are significantly different (P < 0.05).

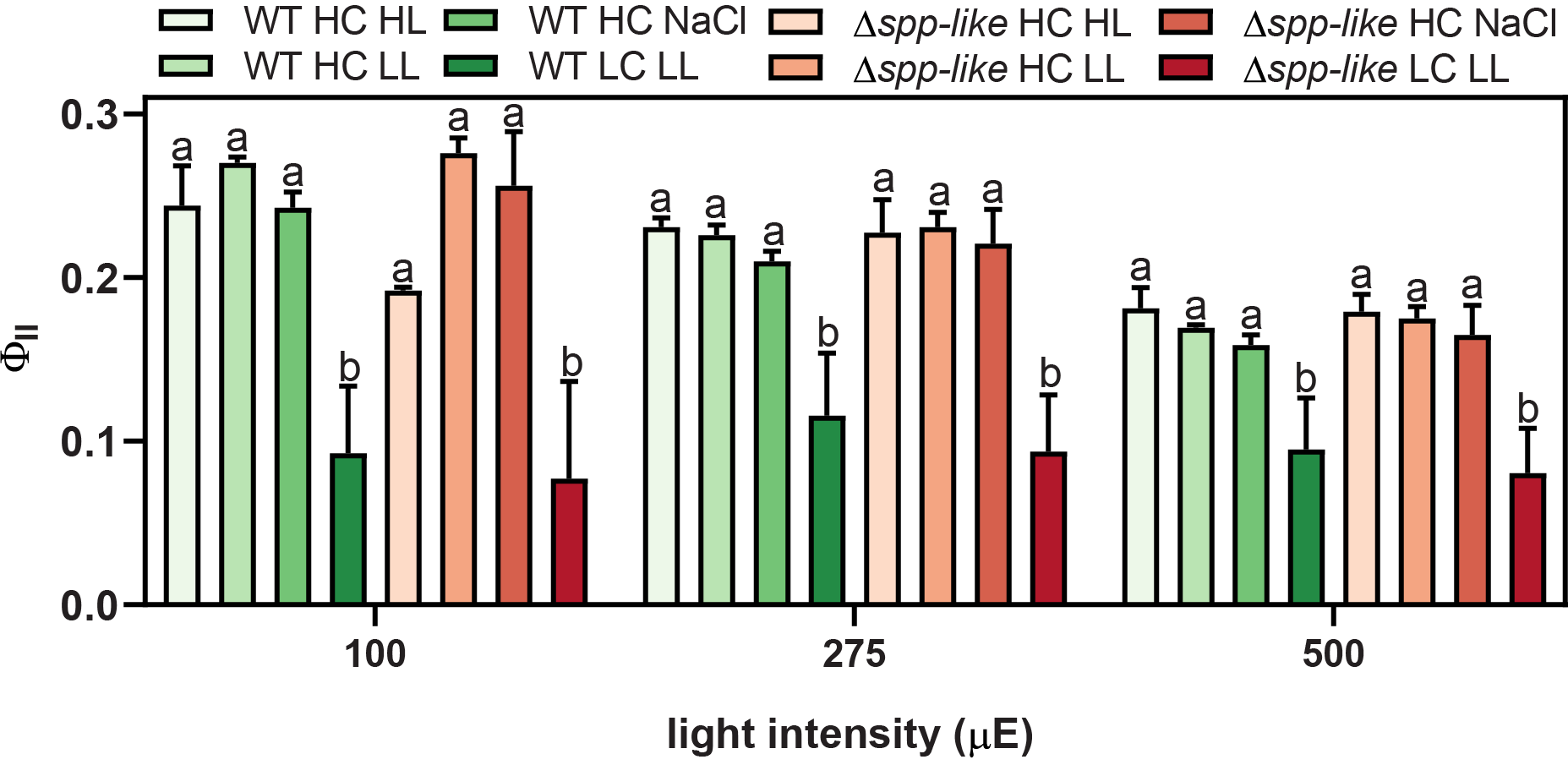

**Figure S11. Photosynthetic performance of the Δ*spp-like* mutant under different growth conditions.** Φ_II_ values were measured at three different light intensities after 24 h of growth under each specific condition. Averages of ≥3 independent biological replicates are shown + SD. Significance was calculated by one-way ANOVA followed by Tukey’s multiple comparison test. Bars labeled with different letters are significantly different (P < 0.05). LC, ambient air CO_2_; HC, 2% CO_2_; HL, 150 μmol photons m^−2^ s^−1;^ LL, 17 μmol photons m^−2^ s^−1^; NaCl, 150 mM NaCl used to induce osmotic stress.

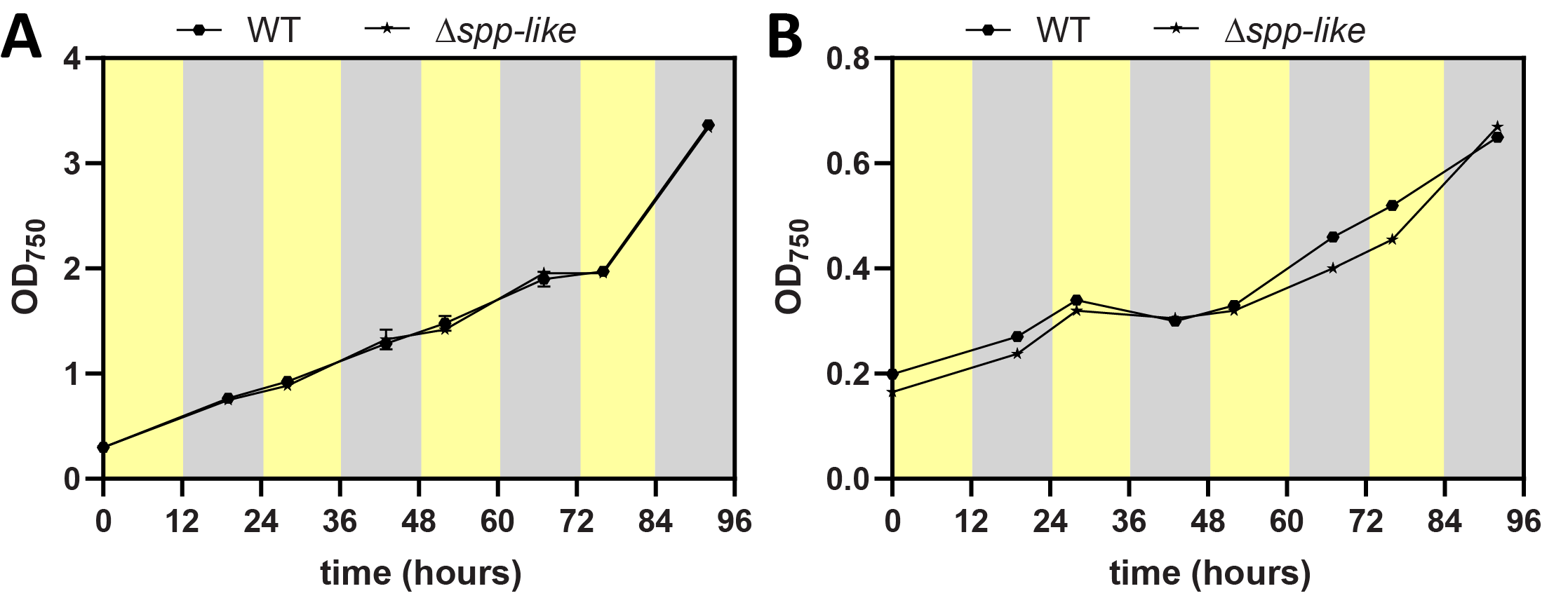

**Figure S12. Light-dark growth curves of the Δ*spp-like* mutant under HC and LC conditions.** Averages of ≥3 independent biological replicates are shown ± SD. LC, ambient air CO_2_; HC, 2% CO_2_. Dark periods are indicated by grey rectangles. Yellow rectangles indicate exposure to a light intensity of 150 μmol photons m^−2^ s^−1^.

3. **Supplementary Tables**

**Table S1.** Protein sequences of orthologs of SPP-like_7942_, SPPu, SPSu and SPPb. *(provided as separate dataset)*

**Table S2.** Cyanobacterial strains used in this study.

**Table S3.** Plasmids used in this study.

**Table S4.** List of the pull-down candidates interacting with SPP-like_7942_ identified by mass spectrometry, including their predicted localization. *(provided as separate dataset)*

**Table S5.** List of significant interactors of SPP-like_7942_ and their predicted localization.

**Table S6.** Enzymes from the SPP-like_7942_ interactome that use phosphorylated metabolite as substrates or products.

**Table S7.** Other relevant proteins from the SPP-like_7942_ interactome with known or unknown functions.

**Table S2.** Cyanobacterial strains used in this study.

| **Strain** | **Relevant genotype and phenotype^a^** | **Plasmids used to generate this strain** | **Source or reference^b^** |
| --- | --- | --- | --- |
| WT | *S. elongatus* PCC 7942 wild type strain. |  | PCC |
| WT/Δ*spp-like* | *S. elongatus* PCC 7942 wild type strain with Δ*spp-like*::*aadA*; Sp^r^ | pSPP-like-KO | This work |
| OE-SPP-like-mNG | *S. elongatus* PCC 7942 wild type strain with P*_trc-RS_*::*spp-like-mNG* integrated at NS1; Sp^r^ | pSPP-like-mNG | This work |
| OE-mNG-SPP-like | *S. elongatus* PCC 7942 wild type strain with P*_trc-RS_*::*mNG*-*spp-like* integrated at NS1; Sp^r^ | pmNG-SPP-like | This work |

^a^Sp^r^, spectinomycin resistance; RS, theophylline-dependent riboswitch; NS1, neutral site 1; mNG, mNeonGreen.

^b^PCC, Pasteur Culture Collection.

Table S3. Plasmids used in this study.

| **Plasmid** | **Relevant genotype and phenotype^a^** | **Source or reference** |
| --- | --- | --- |
| pSPP-like-KO | Plasmid to replace the *spp-like_7942_* gene with Sp^r^ | This work |
| pSPP-like-mNG | Plasmid with P*_trc-RS_*::*spp-like-mNG* for neutral site 1 integration; Sp^r^ | This work |
| pmNG-SPP-like | Plasmid with P*_trc-RS_*::*mNG*-*spp-like* for neutral site 1 integration; Sp^r^ | This work |

^a^Sp^r^, spectinomycin resistance; mNG, mNeonGreen.

**Table S5.** List of significant interactors of SPP-like_7942_ and their predicted localization.

| **Gene name** | **Thylakoid membrane localization^1^** | **CyanoTag localization^2^** |
| --- | --- | --- |
| Synpcc7942_0566 | NA | NA |
| Synpcc7942_1969 | Yes | NA |
| *clpR* | No | D2 |
| *psbD1* | Yes | D1 |
| *psbO* | No | Het: D2, M1 |
| *psaF* | Yes | NA |
| *psaB* | Yes | NA |
| *sqdB* | No | NA |
| *psaD* | Yes | D3 |
| Synpcc7942_1375 | No | NA |
| *prs* | No | D2 |
| *rpaB* | No | NA |
| *nrtA* | Yes | NA |
| *rpsC* | No | NA |
| Synpcc7942_1302 | No | NA |
| *psbA1; psbA2* | Yes | D2 |
| *clpP3* | No | D2 |
| Synpcc7942_1281 | No | NA |
| *clpP1* | No | NA |
| *plmA* | No | NA |
| *yidC* | No | D3 |
| *ftsH* | No | Mixed: M1, P5 |
| Synpcc7942_1634 | No | D2 |
| *ndhI* | No | NA |
| *purF* | No | NA |
| *psaA* | Yes | NA |
| Synpcc7942_0417 | No | NA |
| *petC* | No | D2 |
| *clpC* | No | NA |
| *gpsA* | No | NA |
| *lpxD* | No | NA |
| *serA* | Yes | NA |
| Synpcc7942_2255 | No | NA |
| *accD* | No | NA |
| *aroB* | No | NA |
| *groL* | No | NA |
| *upp* | Yes | NA |
| Synpcc7942_1081 | No | NA |
| *kaiB* | No | D3 |
| *ilvB* | No | NA |
| *rpsS* | No | NA |
| *apcC* | No | NA |
| Synpcc7942_2148 | No | NA |
| Synpcc7942_1065 | No | NA |
| *cobW* | No | NA |
| *metK* | Yes | NA |
| *pgm* | No | NA |
| Synpcc7942_0166 | No | NA |
| *apcE* | Yes | NA |
| 1-cys prx | No | NA |
| *raf1* | No | NA |
| *pyrG* | Yes | NA |
| *recA* | No | NA |
| *plsX* | Yes | NA |
| Synpcc7942_1643 | Yes | NA |
| *synR* | Yes | NA |
| *cpcB1* | No | NA |
| *ccmL* | NA | D1 |
| Synpcc7942_1156 | NA | NA |
| *rpsB* | No | NA |
| *ribD* | NA | LOW |
| *zam* | No | NA |
| *anmK* | No | NA |
| *glpX* | No | D2 |
| *queA* | Yes | NA |
| Synpcc7942_1295 | No | NA |
| *gatA* | No | D2 |
| *thrA* | No | NA |
| *gyrB* | No | NA |
| *hemL* | No | NA |
| *pheT* | No | NA |
| *petH* | No | M1 |
| *atpA* | No | M1 |
| *pilT* | No | NA |
| *flv1* | No | NA |
| *argB* | No | NA |
| *rpsE* | No | NA |
| Synpcc7942_1014 | No | NA |
| *tal* | No | NA |
| Synpcc7942_2306 | Yes | NA |
| Synpcc7942_1429 | No | NA |
| Synpcc7942_1130 | No | NA |
| *gatB* | No | NA |
| *pgk* | No | M1 |
| *pstB* | No | NA |
| *rplX* | No | NA |
| *ama* | NA | NA |
| *glnN* | No | NA |
| *acsA* | Yes | NA |
| *dapB* | No | D3 |
| *ndbA* | Yes | Mixed: P3, M1 |
| *ndhH* | No | NA |
| *fusA* | No | NA |
| *kaiC* | No | Mixed: P3, D2 |
| *era* | No | NA |
| *flv3* | No | Heterogenous: D2, M1 |
| Synpcc7942_2344 | No | NA |
| *dnaN* | No | NA |
| *glyS* | No | NA |
| *rpaA* | No | NA |
| *mtnP* | Yes | NA |
| *rimO* | NA | NA |
| Synpcc7942_0054 | NA | NA |
| Synpcc7942_2499 | No | NA |
| Synpcc7942_0843 | No | NA |
| *ispH* | No | NA |
| *nadB* | NA | NA |
| *murE* | No | NA |
| *infA* | No | NA |
| *hisI* | No | NA |

^1^Thylakoid membrane localization based on (Huokko et al., 2021). NA, not available.

^2^Localization based on (Perrin et al., 2025). D1, diffuse inside the cell, internal to chlorophyll, clearly delineated from chlorophyll; D2, diffuse inside the cell, internal to chlorophyll, no clear delineation; D3, diffuse inside the cell, internal to and overlaps with chlorophyll; M1, surrounds cell body, enriched with chlorophyll with some signal in cell body; P3, punctate/structured, co-incides with chlorophyll; P5, punctate/structured, at cell poles/dividing sites; NA, not available.

**Table S6.** Enzymes from the SPP-like_7942_ interactome that use phosphorylated metabolite as substrates or products.

| **Protein/Gene ID** | **Enzyme** | **Substrate** | **Product** | **Metabolic pathway** |
| --- | --- | --- | --- | --- |
| AroB  Synpcc7942_0525 | 3-dehydroquinate synthase | 3-deoxy-D-*arabino*-hept-2-ulosonate 7-phosphate | 3-dehydroquinate | Shikimate pathway |
| GlpX  Synpcc7942_0505 | D-fructose 1,6-bisphosphatase class 2/sedoheptulose 1,7-bisphosphatase | fructose-1,6-bisphosphate (FBP); sedoheptulose-1,7-bisphosphate | fructose-6-phosphate (F6P); sedoheptulose-7-phosphate | Gluconeogenesis; Calvin cycle |
| Pgm  Synpcc7942_0156 | Phosphoglucomutase  (Reversible reaction) | glucose-1-phosphate (G1P) | glucose-6-phosphate (G6P) | Glycogen synthesis |
| Prs  Synpcc7942_2113 | Ribose-phosphate pyrophosphokinase | ribose-5-phosphate (R5P) | Phosphor-ribosyl pyrophosphate (PRPP) | Nucleotide synthesis |
| PurF  Synpcc7942_0004 | Amidophospho-ribosyltransferase | Phosphor-ribosyl pyrophosphate (PRPP) | Ribosylamine 5-phosphate | Nucleotide synthesis |
| PyrG  Synpcc7942_1954 | CTP synthase | Uridine triphosphate (UTP) | Cytosine triphosphate (CTP) | Nucleotide synthesis |
| SerA  Synpcc7942_1501 | D-3-phosphoglycerate dehydrogenase | 3-phosphoglycerate (3PGA) | 3-phosphohydroxy pyruvate | Amino acid synthesis |
| Upp  Synpcc7942_1715 | Uracil phosphoribosyltransferase | 5-phosphoribosyl-1-pyrophosphate (PRPP) and uracil | Uridine monophosphate (UMP) | Nucleotide synthesis |

**Table S7.** Other relevant proteins from the SPP-like_7942_ interactome with known or unknown functions.

| **Protein/Gene ID** | **Protein description** | **Notes** |
| --- | --- | --- |
| CcmL  Synpcc7942_1422 | Carboxysome shell vertex protein | Essential for carboxysome function |
| GpsA  Synpcc7942_2521 | Nucleotide binding protein, PINc | Function unknown |
| KaiB  Synpcc7942_1217 | Circadian clock oscillator protein | Core structural and regulatory protein in the cyanobacterial circadian clock |
| PlmA  Synpcc7942_0090 | Transcriptional regulator, GntR family | Transcriptional regulator interacting with PipX when PipX is in complex with P_II_ |
| RpaB  Synpcc7942_1453 | Two component transcriptional regulator, winged helix family | Master switch for photosynthesis regulation, stress response, and cell homeostasis and its activity is directly controlled by its phosphorylation state |
| Synpcc7942_1634  Synpcc7942_1634 | Hypothetical protein | Function unknown |
| Synpcc7942_1699  Synpcc7942_1699 | Multidrug and toxic compound transporter | Function unknown |

4. **References**

Cornet, L., Meunier, L., Van Vlierberghe, M., Leonard, R.R., Durieu, B., Lara, Y., Misztak, A., Sirjacobs, D., Javaux, E.J., Philippe, H., Wilmotte, A., Baurain, D. 2018. Consensus assessment of the contamination level of publicly available cyanobacterial genomes. *PLoS One*, **13**(7), e0200323.

Fieulaine, S., Lunn, J.E., Borel, F., Ferrer, J.L. 2005. The structure of a cyanobacterial sucrose-phosphatase reveals the sugar tongs that release free sucrose in the cell. *Plant Cell*, **17**(7), 2049-58.

Huokko, T., Ni, T., Dykes, G.F., Simpson, D.M., Brownridge, P., Conradi, F.D., Beynon, R.J., Nixon, P.J., Mullineaux, C.W., Zhang, P., Liu, L.N. 2021. Probing the biogenesis pathway and dynamics of thylakoid membranes. *Nat Commun*, **12**(1), 3475.

Lazar, C., Gatto, L., Ferro, M., Bruley, C., Burger, T. 2016. Accounting for the multiple natures of missing Values in label-free quantitative proteomics data sets to compare imputation strategies. *J Proteome Res*, **15**(4), 1116-25.

Lin, Y., Wallis, C., Corry, B. 2025. AlphaFold can be used to predict the oligomeric states of proteins. *bioRxiv*, 2025.03.10.642518.

Meng, E.C., Goddard, T.D., Pettersen, E.F., Couch, G.S., Pearson, Z.J., Morris, J.H., Ferrin, T.E. 2023. UCSF ChimeraX: Tools for structure building and analysis. *Protein Sci*, **32**(11), e4792.

Mirdita, M., Schutze, K., Moriwaki, Y., Heo, L., Ovchinnikov, S., Steinegger, M. 2022. ColabFold: making protein folding accessible to all. *Nat Methods*, **19**(6), 679-682.

Old, W.M., Meyer-Arendt, K., Aveline-Wolf, L., Pierce, K.G., Mendoza, A., Sevinsky, J.R., Resing, K.A., Ahn, N.G. 2005. Comparison of label-free methods for quantifying human proteins by shotgun proteomics. *Mol Cell Proteomics*, **4**(10), 1487-502.

Perez-Riverol, Y., Bai, J., Bandla, C., Garcia-Seisdedos, D., Hewapathirana, S., Kamatchinathan, S., Kundu, D.J., Prakash, A., Frericks-Zipper, A., Eisenacher, M., Walzer, M., Wang, S., Brazma, A., Vizcaino, J.A. 2022. The PRIDE database resources in 2022: a hub for mass spectrometry-based proteomics evidences. *Nucleic Acids Res*, **50**(D1), D543-D552.

Perrin, A.J., Dowson, M., Davis, K., Nam, O., Dowle, A.A., Calder, G., Springthorpe, V.J., Zhao, G., Mackinder, L.C.M. 2025. CyanoTag: Discovery of protein function facilitated by high-throughput endogenous tagging in a photosynthetic prokaryote. *Sci Adv*, **11**(6), eadp6599.

Santos-Merino, M., Torrado, A., Davis, G.A., Rottig, A., Bibby, T.S., Kramer, D.M., Ducat, D.C. 2021. Improved photosynthetic capacity and photosystem I oxidation via heterologous metabolism engineering in cyanobacteria. *Proc Natl Acad Sci U S A*, **118**(11).

Teikari, J., Baunach, M., Dittmann, E. 2022. Cyanobacterial genome sequencing, annotation, and bioinformatics. *Methods Mol Biol*, **2489**, 269-287.

Trifinopoulos, J., Nguyen, L.T., von Haeseler, A., Minh, B.Q. 2016. W-IQ-TREE: a fast online phylogenetic tool for maximum likelihood analysis. *Nucleic Acids Res*, **44**(W1), W232-5.

Tyanova, S., Temu, T., Cox, J. 2016. The MaxQuant computational platform for mass spectrometry-based shotgun proteomics. *Nat Protoc*, **11**(12), 2301-2319.
